# Single-nucleus transcriptional profiling of GAD2-positive neurons from mouse lateral habenula reveals distinct expression of neurotransmission- and depression-related genes

**DOI:** 10.1101/2023.01.09.523312

**Authors:** Matthew V. Green, David A. Gallegos, Jane-Valeriane Boua, Luke C. Bartelt, Arthy Narayanan, Anne E. West

**Affiliations:** Department of Neurobiology, Duke University, Durham NC 27710

## Abstract

Glutamatergic projection neurons of the lateral habenula (LHb) drive behavioral state modulation by regulating the activity of midbrain monoaminergic neurons. Identifying circuit mechanisms that modulate LHb output is of interest for understanding control of motivated behaviors. A small population of neurons within the medial subnucleus of the mouse LHb express the GABAergic synthesizing enzyme GAD2, and they can inhibit nearby LHb projection neurons; however, these neurons lack markers of classic inhibitory interneurons and they co-express the vesicular glutamate transporter VGLUT2. To determine the molecular phenotype of these neurons, we genetically tagged the nuclei of GAD2-positive cells and used fluorescence-activated nuclear sorting to isolate and enrich these nuclei for single nuclear RNA sequencing (FANS-snRNAseq). Our data confirm that GAD2+/VGLUT2+ neurons intrinsic to the LHb co-express markers of both glutamatergic and GABAergic transmission and that they are transcriptionally distinct from either GABAergic interneurons or habenular glutamatergic neurons. We identify gene expression programs within these cells that show sex-specific differences in expression and that are implicated in major depressive disorder (MDD), which has been linked to LHb hyperactivity. Finally, we identify the *Ntng2* gene encoding the cell adhesion protein Netrin-G2 as a marker of LHb GAD2+/VGLUT+ neurons and a gene product that may contribute to their target projections. These data show the value of using genetic enrichment of rare cell types for transcriptome studies, and they advance understanding of the molecular composition of a functionally important class of GAD2+ neurons in the LHb.

## INTRODUCTION

The habenula is a hub for a complex network of anatomical connections that link the limbic forebrain with modulation of midbrain monoaminergic nuclei. The integration of forebrain inputs in the habenula, and the consequent effects on the activity of dopaminergic, serotonergic, and noradrenergic neurons, effect behavioral state modulation of motivation, reward processing, and social approach/avoidance behaviors (Hikosaka, 2010). The habenula is part of the epithalamus and is comprised of two major subregions: the medial habenula (MHb) and lateral habenula (LHb), which are defined by distinct anatomical connectivity and unique behavioral functions (Hikosaka, 2010). The LHb receives inputs from hypothalamus, cortex, ventral tegmental area (VTA), dorsal raphe nuclei (DRN), and locus coeruleus (LC), and then sends both direct and indirect projects to modulate the firing of monoaminergic neurons in VTA, DRN, and LC affecting a wide range of functions (Hu et al., 2020). Dysfunction of the LHb is thought to contribute to psychiatric disorders that are associated with dysregulation of motivation and reward, such as drug addiction and depression (Hu et al., 2020).

The LHb is composed predominantly of glutamatergic neurons that project to downstream target regions in the midbrain (Hashikawa et al., 2020; Jhou et al., 2009; Wallace et al., 2020). Cellular and synaptic mechanisms that promote activity of these excitatory projections have been associated with depressive-like behaviors in animal models (Li et al., 2011; Shabel et al., 2014; Tchenio et al., 2017; Yang et al., 2018), whereas inhibition of LHb activity has been suggested as a potential therapeutic option for major depressive disorder (MDD) (Sartorius et al., 2010). These observations have driven substantial interest in identifying sources of circuit inhibition that can reduce the firing of LHb projection neurons. GABAergic afferents that project to the LHb have been traced from multiple brain regions including the entopeduncular nucleus, the ventral pallidum, the VTA, the hypothalamus, and the dorsomedial nucleus of the thalamus (Webster et al., 2021).

Inhibition of LHb efferents could also be mediated by local GABAergic neurons. A small population of GABA+ neurons that co-express the GABAergic interneuron marker parvalbumin together with the vesicular GABA transporter (VGAT) are found in far lateral regions of LHb, and stimulation of these neurons can induce IPSCs in neighboring LHb neurons (Webster et al., 2020). An independent population of neurons expressing the GABA synthesizing enzyme GAD2 are found in the medial subnucleus of the LHb, however these cells do not express VGAT, GAD1, or other canonical interneuron markers and they co-express the vesicular glutamate transporter VGLUT2 (Quina et al., 2020). Using *Gad2*-Cre mice in combination with Cre-dependent AAV-ChR2 expression in the LHb, one group found that optogenetic stimulation of these neurons resulted in inhibitory currents recorded from locally connected neurons within the LHb (Flanigan et al., 2020). However, using a similar strategy, another group recorded excitatory currents when they followed projections from these LHb GAD2+ neurons to their targets in the mesopontine tegmentum (Quina et al., 2020). The molecular mechanisms that allow this GAD2+ cell population to have properties of both local inhibitory interneurons and excitatory projection neurons has remained unknown.

Single cell sequencing has revolutionized neuronal classification by reducing traditional reliance on categorical marker genes and allowing for the holistic analysis of gene sets that define cellular function (Huang & Paul, 2019). Recent single cell sequencing papers have characterized cell type diversity within the medial and lateral habenula, yielding important insights into the relevant gene expression programs that characterize downstream projection patterns (Cerniauskas et al., 2019; Hashikawa et al., 2020; Pandey et al., 2018; Wallace et al., 2020). Although a small number of *Gad2* expressing cells were detected in these studies, there were insufficient numbers to drive these neurons into a cluster of their own, and thus the transcriptome of this population remains uncharacterized. To overcome this limitation, we and others have been using transgenes to enrich single cells (Munoz-Manchado et al., 2018) or single nuclei (Gallegos et al., 2022) of rare cell types by fluorescence activated sorting (FACS/FANS) from heterogeneous brain tissues prior to sequencing. Here, we provide and analyze a comprehensive single nuclear RNA-seq (snRNA-seq) dataset of nuclei from GAD2+ neurons enriched from the LHb. By integrating our Gad2-enriched dataset with previous total scRNA-seq data from the habenula, we describe the expression of key genes that define the transcriptional profiles of LHb GAD2+ neurons compared with other LHb neurons, and we identify the axon guidance gene *Ntng2* (encoding the axon guidance molecule NetrinG2) as a marker that differentiates the population of *Gad2/Slc17a6* co-expressing neurons in the LHb.

## RESULTS

### Single-nuclear gene expression profiles define distinct populations of LHb Gad2+ neurons

To obtain comprehensive transcriptional profiles of LHb *Gad2*-expressing cells, we genetically tagged and enriched for the nuclei of these neurons prior to performing snRNA-seq. We crossed mice expressing a Cre-inducible, GFP-tagged nuclear envelope protein (ROSA-LSL-Sun1-sfGFP) (Mo et al., 2015) with mice expressing Cre recombinase knocked into the *Gad2* locus (*Gad2*-IRES-Cre)(Taniguchi et al., 2011) to label the nuclei of *Gad2*+ cells throughout the brain (**Fig. S1A,B)**. We then microdissected the LHb from adult male and female mice, harvested all nuclei and then performed isolation of GFP+ nuclei by FACS-assisted nuclear sorting (FANS) followed by single-nuclear RNA sequencing (snRNA-seq) on the 10X Genomics platform (**Fig. 1A; Fig. S1C**).

**Figure 1:**
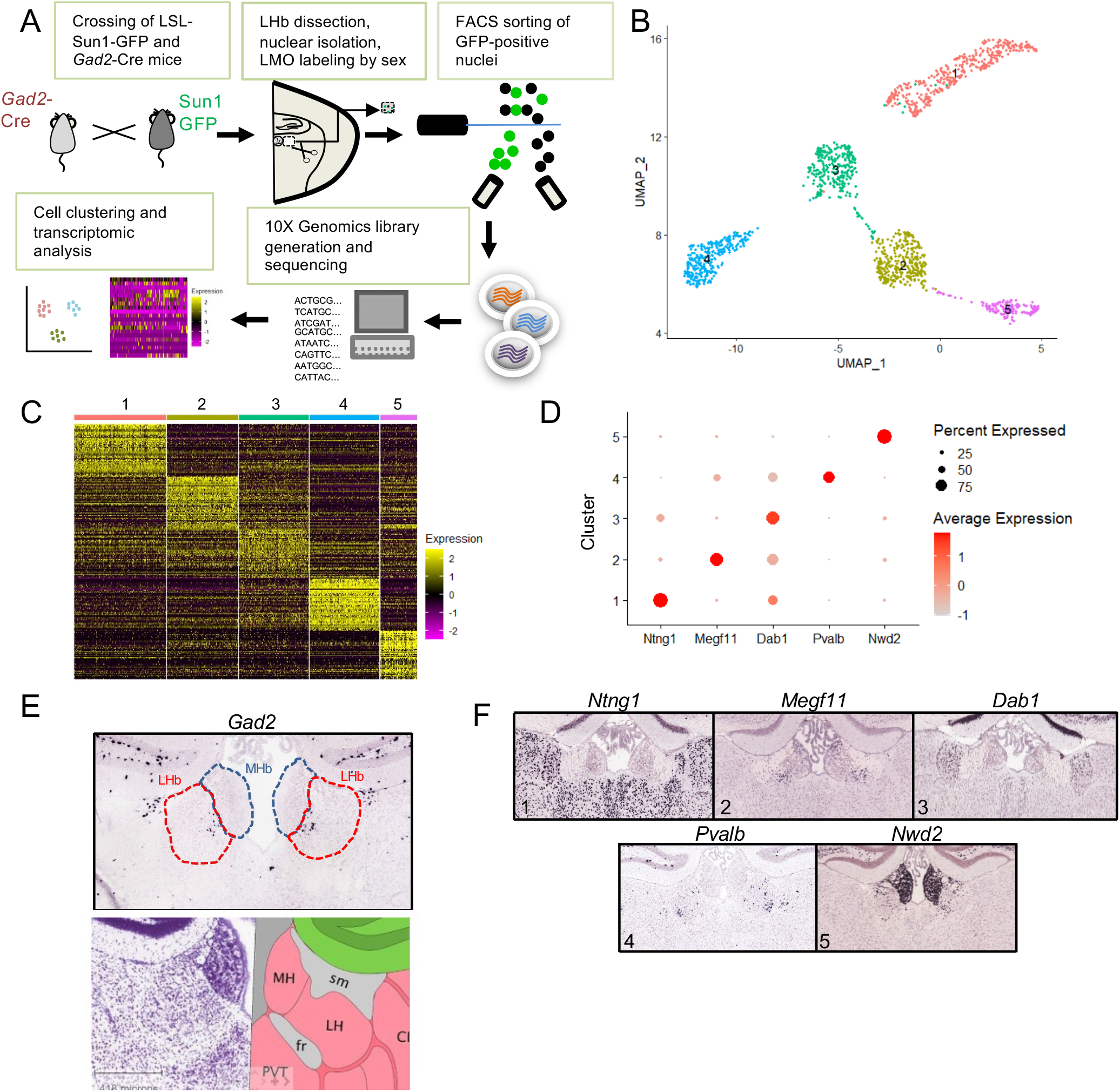
Isolation and snRNA-seq analysis of Gad2+ neurons from the LHb. **(A)** Schematic of the approach designed to enrich for Gad2+ neurons within the LHb and subsequent snRNA-seq. **(B)** UMAP clustering of the 5 neuronal clusters retrieved from snRNA-seq, filtered by expression of *Rbfox3*. **(C)** Heatmap showing differential expression (DE) of top 50 genes per cluster using a non-parametric Wilcoxon Rank Sum test, adjusted p-value ≤ 10e^-6^ and log_2_foldchange ≥ |0.25|. Columns represent each cluster with sub columns representing single cells. Each row represents a different gene. **(D)** Dot plot showing scaled expression of representative top DE genes per cluster. **(E)** Spatial expression from the Allen Institute Brain Atlas showing *Gad2* expression enriched in the medial portion of the LHb. A reference map is shown for orientation. **(F)** Representative top DE genes per cluster to highlight the respective anatomical origins.

We recovered transcriptomes from 3,651 GFP+ cells and performed principal component analysis to reduce the dimensionality of the data and allow for subsequent clustering analysis and visualization using the UMAP algorithm (**Fig. S2**). This method identified 11 clusters of cells. Almost all the clusters contained some nuclei with *Gad2* mRNA (**Fig. S2C**) and variation in *Gad2* expression between cells in a cluster could reflect RNA dropout. However, when we assessed the expression of known cell-type marker genes within these clusters we found that many were comprised of non-neuronal cells (**Fig. S2B**). This may reflect expression of the *Gad2*-Cre transgene in some non-neuronal cells, as we did observe some GFP expression in a population of NeuN-cells in the habenula and surrounding structures **(Fig. S1B)**. Because we wanted to focus our analysis on neurons, we filtered clusters for the expression of the neuronal marker *Rbfox3* (**Fig. S2D**). This resulted in five clusters of 1491 cells (**Fig. 1B**) with similar QC features between clusters **(Fig S3)**. The expression of activity-inducible genes was uniform across the neuronal clusters supporting that these neurons clustered by cell type rather than activity state (**Fig. S4**). indicating that the sorting had successfully enriched for nuclei with *Gad2*-Cre driven Sun1-GFP

We assessed the most differentially expressed (DE) genes between clusters (**Fig. 1C; Table S1**), and then used a subset of these highly differentially expressed genes (**Fig. 1D**) in combination with the Allen Brain Atlas *in situ* hybridization data to verify the original anatomical origins of these neuronal clusters (**Fig. 1E)**. Clusters 2 and 4 are marked by high expression of *Megf11* and *Pvalb* respectively, both of which have expression patterns in the LHb that have similarities to the pattern of *Gad2* in this brain region. *Pvalb*+ neurons are also found in the dorsal thalamus, which immediately borders the LHb on the lateral side. Not surprisingly, given the small size of the LHb and the sparse but broad distribution of *Gad2*+ cells in this region of the brain, some of the clusters appear to have originated from regions outside of but in in close apposition to the LHb. Markers of clusters 1 (*Ntng1*) and 3 (*Dab1*) are enriched in the dorsal thalamus, whereas cluster 5 expressed high levels of *Nwd2*, which is highly expressed in the medial habenula (MHb).

Of the five neuronal clusters, clusters 2, 3, and 4 expressed the highest levels of *Gad2* (**Fig. 2A**). Given our marker gene evidence (**Fig. 1F**) that clusters 2 and 4 are the most likely to represent *Gad2*+ neurons arising from the LHb, we focused on comparing gene expression between these two clusters. Unlike canonical cortical GABAergic interneurons (Paul et al., 2017), *Gad2*-expressing cells in the medial LHb are known for their co-expression of *Slc17a6* encoding the excitatory glutamate vesicular transporter VGLUT2 (Quina et al., 2020). We saw that cluster 2 but not cluster 4 expressed *Slc17a6* as well as *Gad2* (**Fig. 2B-C**). Because these clusters represent groups of nuclei, we graphed the levels of *Gad2* expression against *Slc17a6* expression for each single nucleus in each of the five clusters (**Fig. S5**). Whereas most nuclei in clusters 1 and 5 showed *Slc17a6* expression and most nuclei in clusters 3 and 4 showed *Gad2* expression, only cluster 2 had a significant number of cells that showed expression of both transporter genes.

**Figure 2:**
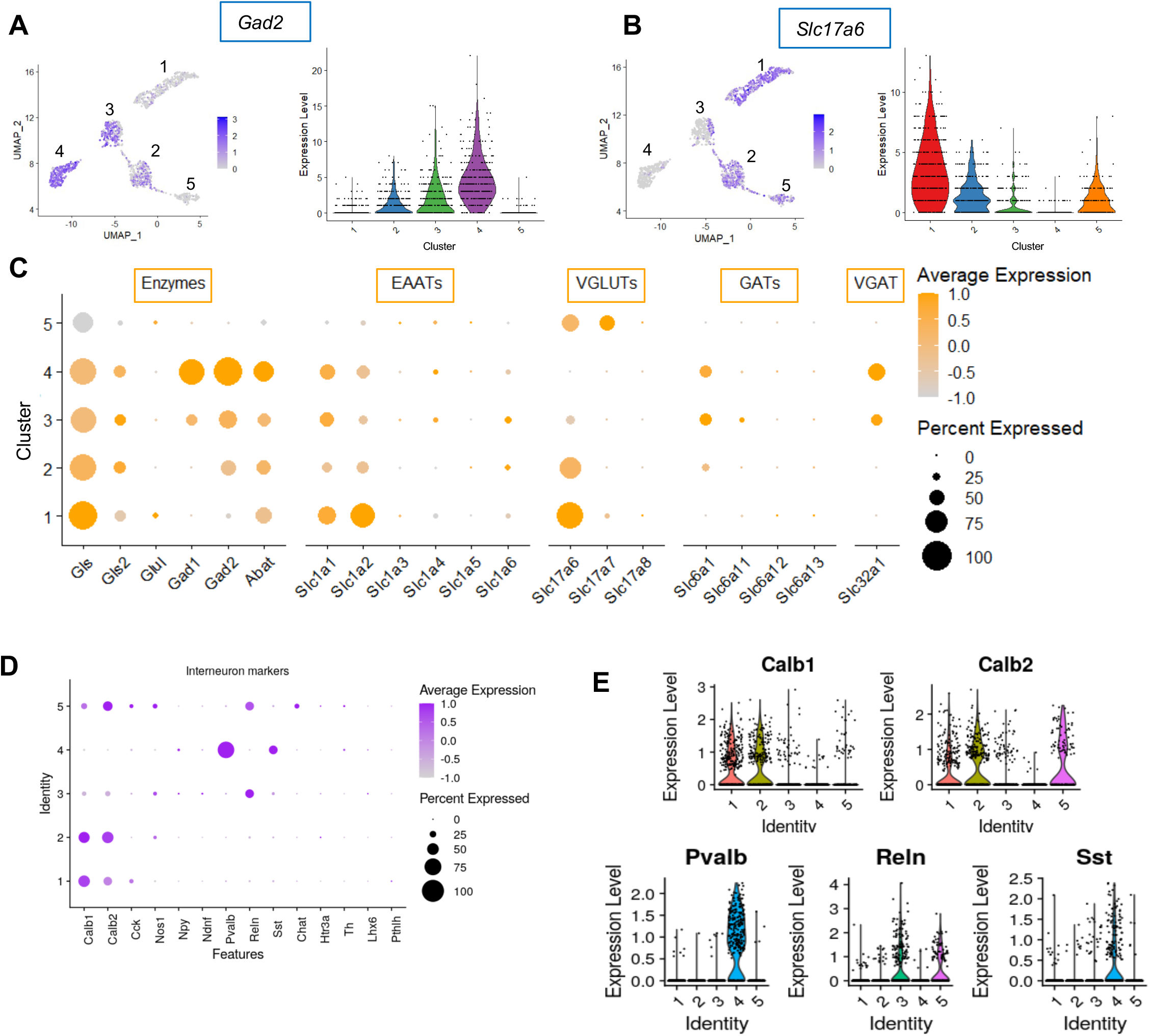
Gad2+ LHb neurons in cluster 2 express both GABAergic and glutamatergic transmitter features. **(A**,**B)** Feature plots showing scaled expression and violin plots showing raw counts of *Gad2* **(A)** and *Slc17a6* **(B)** expression within the five neuronal Gad2+ clusters. **(C)** Dot plot showing scaled expression of genes related to synthesis, degradation, release, and reuptake of glutamate and GABA across the five neuronal Gad2+ clusters. **(D)** Dot plot and (**E)** violin plots showing scaled expression of GABAergic interneuron-associated markers in the five neuronal Gad2+ clusters.

We next assessed the distribution of genes that regulate the synthesis, release, reuptake, and degradation of GABA. Cluster 4 but not cluster 2 neurons expressed *Gad1*, encoding the GABA synthesizing enzyme GAD67 **(Fig 2C)**. Further consistent with cluster 4 representing a subtype of GABAergic inhibitory interneurons, we found that this was the only cluster strongly expressing the interneuron markers *Pvalb* and *Sst* **(Fig. 2D,E)** (Yuste et al., 2020). By contrast, we observed little expression of any of the usual interneuron class markers in cluster 2, though these cells did express *Calb1*, encoding Calbindin and *Calb2*, encoding Calretinin **(Fig. 2D,E)**. Consistent with prior studies of the *Gad2*+ neurons in the medial subnucleus of the LHb, we found that neurons in cluster 2 did not express *Slc32a1*, encoding the vesicular GABA transporter VGAT, whereas this gene was strongly expressed in the *Gad2+* neurons of cluster 4. Interestingly however, neurons in cluster 2 were seen to express *Slc6a1*, the gene encoding the plasma membrane GABA transporter GAT1 **(Fig. 2C**). This is important because GAT1 function has been identified as a possible mechanism for release of GABA by transporter reversal under membrane depolarized conditions even from neurons that lack VGAT to support vesicular GABA release (Attwell et al., 1993; Cammack et al., 1994).

To validate whether *Gad2*+ cells in the medial subnucleus of the LHb simultaneously co-express both the glutamate transporter *Slc17a6* and the plasmid membrane GABA transporter *Slc6a1*, we used RNAscope fluorescent *in situ* hybridization (FISH) to quantify the expression and colocalization of these markers on brain sections (**Fig. 3)**. *Slc17a6* was expressed in a large percentage of cells across the MHb and LHb, consistent with the evidence that most neurons in this region are glutamatergic (**Fig. S6; Fig. 3B, F**). *Slc6a1* was found in both MHb and LHb, though it was expressed at higher levels and in a greater percentage of cells in the LHb compared with the MHb (**Fig. S6; Fig. 3C, F**). Within the LHb, *Slc17a6* and *Slc6a1* were not limited to *Gad2*+ cells, but either one or both were co-expressed with *Gad2* in the majority of *Gad2*+ cells. (**Fig. 3D,E,G**). Taken together with our snRNA-seq analysis, these data confirm that cluster 2 likely represents the medial LHb *Gad2*+ cell cluster of functional interest for its coexpression of GABAergic and glutamatergic transmitter genes.

**Figure 3:**
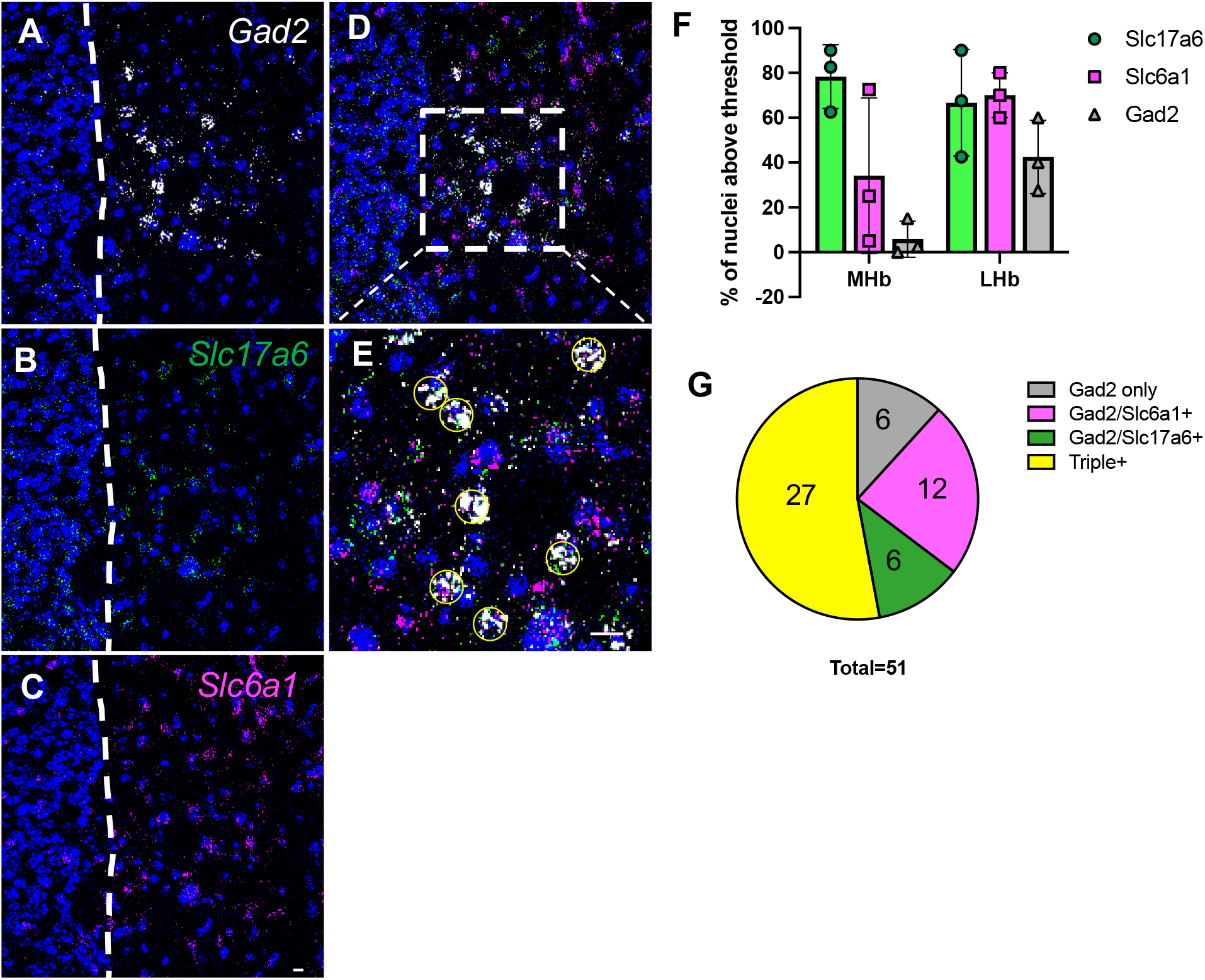
Most *Gad2*+ neurons in the medial subnucleus of the LHb co-express *Slc17a6* and/or *Slc6a1*. **(A-C)** Representative RNAscope ISH images of *Gad2* **(A)**, *Slc17a6* **(B)**, and *Slc6a1* **(C)** in the habenula. DAPI, nuclei (blue). Dotted white line, boundary between the MHb (left) and LHb (right). **(D)** Overlap of *Gad2* with *Slc17a6* and *Slc6a1*. Dotted white box is centered on the medial subnucleus of the LHb and expanded in **(E)**. Yellow circles show *Gad2*+ cells evaluated for overlap. **(F)** Quantification of the percent of positive nuclei for each individual brain section from thresholding in **Fig. S6**. 40 cells/image/channel from 3 brains. **(G)** Percentage of the total of all *Gad2*+ positive nuclei from images of 3 brains that co-express *Slc17a6, Slc6a1*, or both. A total of 51 cells were quantified on 3 brains. Scale bar: 10 μm.

### Gad2+/Slc17a6+ LHb neurons are distinguished by distinct neurotransmitter gene expression profiles

We further investigated the transcriptional profile of cluster 2 by looking for the differential expression of key gene families that could provide insight into their function. First, we identified the strongest DE genes between cluster 2 and the other four *Gad2*+ neuron clusters from our sample using a Wilcoxon Rank Sum test (**Fig. 4A, Table S2**). GO analysis of the DE genes identified glutamatergic synapse genes and other components of chemical synaptic transmission as defining categories (**Fig. 4B; Table S3**). In cases where the same GO category was significant in both the genes that were significantly enriched and de-enriched in cluster 2, this is because different members of the same gene family were expressed in the different cell clusters. For example, in the molecular function (MF) category glutamate receptor activity (GO:0008066), *Gria1, Gria2*, and *Grik4* were higher in cluster 2 relative to the other clusters, whereas *Gria4, Grik2*, and *Grik3* were lower. Indeed, cluster 2 expressed a unique profile of genes for glutamatergic receptors in all of the families, notably including low expression of genes for NMDA-type glutamate receptor subunits as well differential expression of subunits for AMPA-type, metabotropic, and kainate-type glutamate receptors (**Fig. 4C**). Among the GABA receptor genes, neither clusters 2 nor 4 showed especially high expression especially compared with cluster 1, but cluster 2 displayed higher expression of *Gabrg1* and cluster 4 had highest expression of *Gabra3*, again suggesting distinctions in the composition of GABA receptors among different classes of *Gad2*+ neurons (**Fig. 4D**).

**Figure 4:**
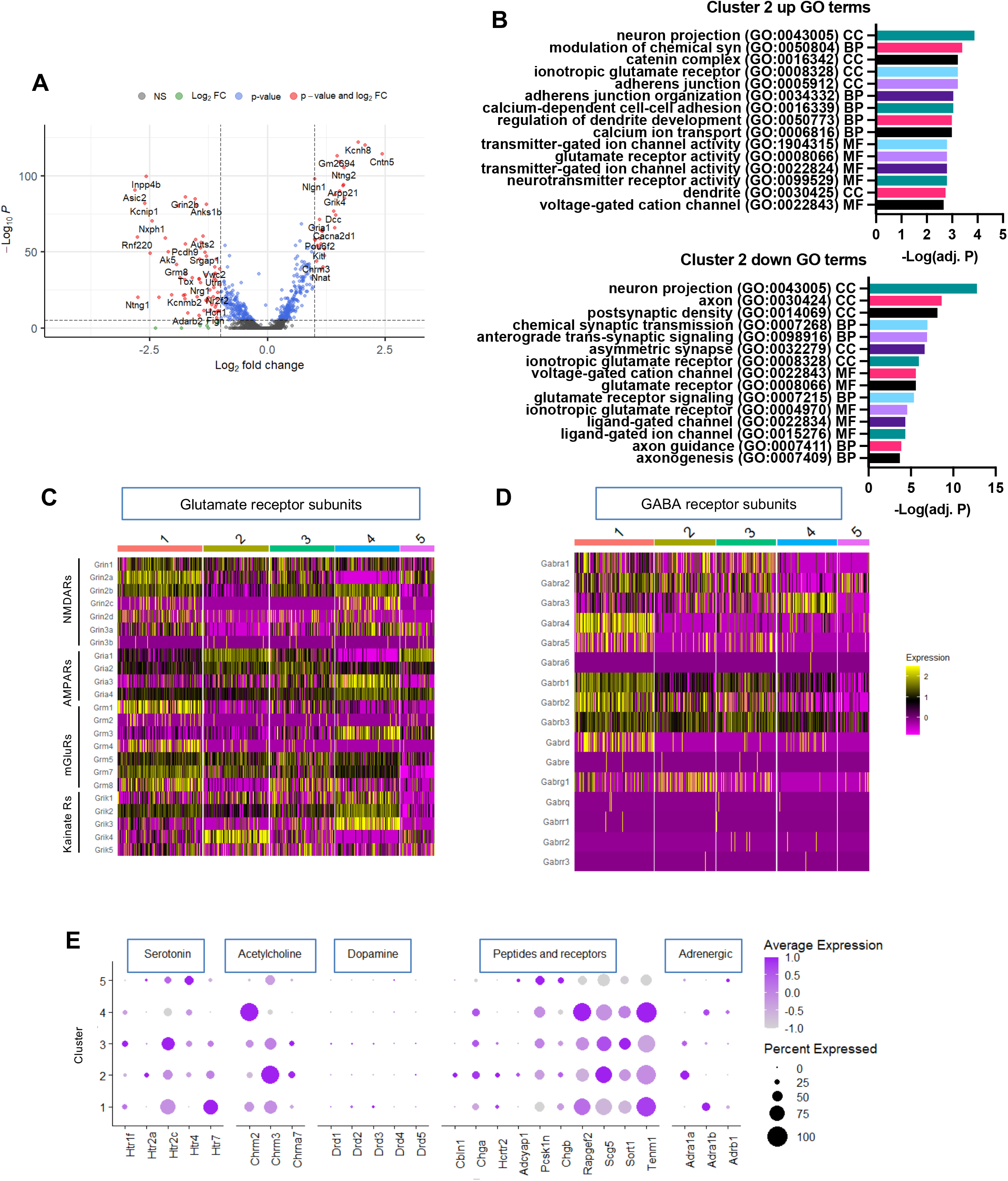
Neurotransmitter and neuromodulator receptor profiles distinguish different Gad2+ neuronal populations in the LHb. **(A)** Volcano plot highlighting top DE genes for cluster 2 compared to the other 4 neuronal clusters in the dataset using a non-parametric Wilcoxon Rank Sum test. Dashed lines indicate cut-offs in the horizontal and vertical direction for log2foldchange ≥ |0.25| and adjusted p-value ≤ 10e^-6^, respectively. **(B)** GO analysis summary showing the top 5 GO terms in each category (molecular function, MF, biological process, BP, and cellular compartment, CC) for the up and down-regulated DE genes, as determined from **A**. Heatmap displaying expression of glutamate receptor subunits and **(D)** GABA receptor subunits per cluster. Each sub column within a cluster represents a single nucleus. Scaled expression is shown. **(E)** Dot plot showing expression of the following gene categories within each of the five Gad2+ neuronal clusters: serotonin receptors, acetylcholine receptors, dopamine receptors, peptides and their receptors, and adrenergic receptors.

Next, because the LHb is implicated in behavioral state integration, we examined whether cluster 2 was marked by distinctive expression of genes involved in neuromodulation. All our *Gad2*+ clusters strongly expressed genes encoding serotonin and acetylcholine receptors and they weakly expressed noradrenergic receptor subunits; however, we saw minimal expression of dopamine receptors in any of the clusters (**Fig. 4E)**. The most strikingly differential expression between clusters is among the acetylcholine receptors, with *Chrm3* highly expressed in cluster 2 versus *Chrm2* in cluster 4. *Chrm3* was identified in a previous scRNA-seq study of the habenula as a marker of cells in both the lateral oval as well as the medial subdivisions of the LHb (Wallace et al., 2020). Given that *Gad2* expression is restricted to the medial subdivision, our findings suggest these are distinct *Chrm3+* cell types in the two regions.

### Gad2/Slc17a6 double positive LHb neurons are transcriptionally distinct from other Slc6a1+ LHb neurons

Our enrichment strategy allowed us to sequence many *Gad2*+ neurons from the LHb, providing a robust transcriptional analysis of these rare neurons. To compare the programs of gene expression we observed in our *Gad2*+ clusters against those of *Gad2*-neurons of the LHb, we integrated our dataset with a previously published scRNA-seq dataset from the MHb and LHb (Hashikawa et al., 2020). Clustering of the integrated datasets confirmed our identification of microglia, astrocytes, oligodendrocytes, and endothelial cells from our preliminary clusters (**Fig. S2, S7**). The co-clustering of these non-neuronal cell types between the two datasets demonstrates the feasibility of integrating scRNA- and snRNA-seq data from independent experiments performed by different laboratories. After filtering the integrated dataset for only *Rbfox3+* neuron-containing clusters, we obtained a total of 12 clusters. We call these clusters I1-I12 (I for “Integrated”) to distinguish them from the 5 *Gad2+* clusters in our data from **Fig. 1**. This clustering supports our conclusion that cluster 5 of our *Gad2+* neurons (**Fig. 1B**) contained cells from the MHb, as these neurons colocalized in clusters I2, I4, and I7 with MHb cells from the Hashikawa et al. (2020) dataset (**Fig. 5A,B; S8**). As expected, the perihabenular *Gad2*+ cells from our clusters 1 and 3 failed to overlap with the habenular clusters. The highly *Gad2*+ cells in cluster 4 that we suggested were canonical interneurons **(Fig. 2)** most closely cluster with peri-habenular neurons in I9, suggesting either that they may not arise from the LHb, or that they may be too small in number in the larger dataset to permit them to cluster as LHb interneurons.

**Figure 5:**
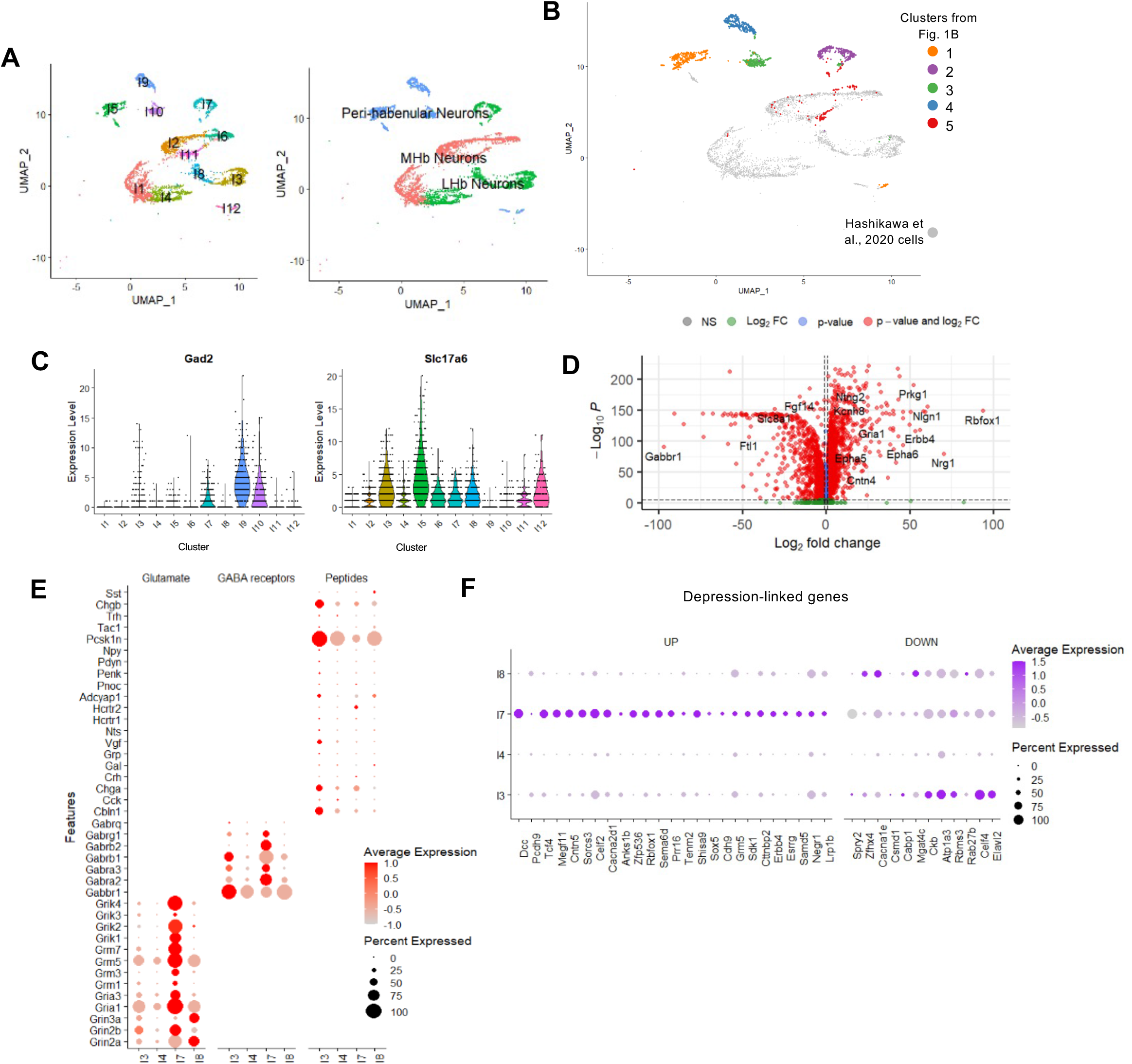
*Gad2/Slc6a17* dual-positive neurons of the LHb are transcriptomically distinct from other LHb neurons. **(A)** UMAP clustering of all neurons after integration with Hashikawa et al., 2020 dataset. **(B)** UMAP clustering of all neuron clusters with neurons from our dataset highlighted according to original cluster number from Figure 1B versus neurons from Hashikawa et al., 2020 data set (grey). **(C)** Violin plot of expression of *Gad2* and *Slc17a6* across the 12 neuronal clusters in the integrated dataset. **(D)** Volcano plot highlighting top DE genes between the LHb clusters. DE was run using a non-parametric Wilcoxon Rank Sum test. Dashed lines indicate cut-offs in the horizontal and vertical direction for log2foldchange ≥ |0.25| and adjusted p-value ≤ 10e^-32^, respectively. **(E)** Dot plot showing expression of glutamate receptor subunits, GABA receptor subunits, and peptides between cluster I7 compared with I3, I4, and I8. Scaled expression is shown. **(F)** Dot plot showing the enriched (UP) and de-enriched (DOWN) genes in cluster I7 compared with clusters I3, I4, and I8 from cross-reference to a meta-analysis of 269 depression-linked genes. Scaled expression is shown.

Interestingly, the cells we identified as the *Gad2/Slc17a6* dual-expressing cluster 2 from **Fig. 1B** clustered on their own in the integrated dataset as integrated cluster I7 (**Fig 5A,B**). Furthermore, cluster I7 was the only cluster that showed significant co-expression of *Gad2* and *Slc17a6* in the entire integrated dataset (**Fig. 5C**). These data reinforce the power of our enrichment strategy to determine the transcriptome of a rare cell population that was missed by clustering of cells from the total habenula.

We ran differential expression analysis between cluster I7 and the three clusters from the Hashikawa et al., 2020 dataset that contain LHb neurons (clusters I3, I4 and I8; **Fig. 5A, Fig. S7-S8, Table S2)** to determine which genes most strongly distinguish the *Gad2/Slc17a6* double-positive neurons from *Gad2*-neurons in the LHb (**Fig. 5D**). Prior studies have shown that *Gad2*+ cells of the medial subnucleus express orexin receptors (*Hcrtr2*) that mediate the modulatory effects of orexin on aggression (Flanigan et al., 2020). We confirmed the preferential expression of the orexin receptor *Hcrtr2* in cluster I7 relative to other LHb neurons (**Fig. 5E**), though we did not find that these neurons were particularly enriched for other peptide receptors (Flanigan et al., 2020). We confirmed the enrichment of *Chrm3* in cluster I7 relative to the *Gad2*-LHb neurons and noted a relative paucity of neuromodulator receptors in any of the LHb neurons except the *Gad2/Slc17a6* double positive cells in cluster I7 **(Fig. S9**).

Finally, because the LHb has been implicated in MDD and antidepressant action (Gold & Kadriu, 2019; Hu et al., 2020; Li et al., 2011; Shabel et al., 2014), we sought to determine whether there were any significant DE genes preferentially expressed in cluster I7 versus other neurons in the LHb that were genetically associated with MDD. This could implicate the function of Gad2+ neurons in emotional regulation. A recent meta-analysis (Howard et al., 2019) yielded a list of 269 genes with significant genetic risk for MDD. We first cross-referenced this list against the set of genes that were preferentially expressed in cluster I7 *Gad2/Slc17a6*+ neurons relative to *Gad2*-LHb neurons from clusters I3, I4, and I8 (**Table S4)**. We identified 25 MDD-associated genes that were preferentially expressed in cluster I7 relative to other LHb neurons (**Fig. 5F)**. For comparison, only 12 of the 269 genes were more strongly enriched in any of the three other LHb clusters compared with I7. Of the I7-enriched genes, 10 were also significantly enriched in cluster 2 relative to other Gad2+ neurons from LHb (*Dcc, Tcf4, Megf11, Cacna2d1, Sorcs3, Erbb4, Chd9, Sema6d, Zfp536*, and *Sdk1*25) (**Fig S10**). Whether these genes have specific functions in Gad2+ neurons of the LHb that underlie the role of this brain region in MDD will be of interest to test in the future.

### Sex-specific differences in LHb Gad2+ neuron gene expression

A prior study has shown that *Gad2/Slc17a6* double-positive LHb neurons express estrogen receptors and suggested that gonadal steroid-dependent modulation of these neurons might contribute to sex-specific and behaviorally relevant regulation of LHb activity (Zhang et al., 2018). To determine whether *Gad2*+ LHb neurons show sex-specific differences in gene expression, we tagged the nuclei harvested from either male or female mice with distinct lipid-modified oligonucleotide barcodes (McGinnis et al., 2019). After pooling for sequencing, we used to the barcodes to deconvolve *Gad2*+ neurons that came from male or female mice. We confirmed that the Y-chromosome genes *Eif2s3y, Uty, Kdm5d*, and *Ddx3y* were expressed only in nuclei marked by the barcode we added to nuclei of male mice, whereas *Xist* and *Tsix*, which mediate X-chromosome inactivation, were found exclusively in nuclei containing the female barcode (**Fig. 6A**). These data show that we can use LMO barcoding to retrospectively subset nuclei as originating from male or female mice for sex-specific comparison of gene expression. After sex classification and filtering for expression of the neuronal marker *Rbfox3*, as described in the methods section, we unambiguously identified 816 male and 107 female nuclei.

**Figure 6:**
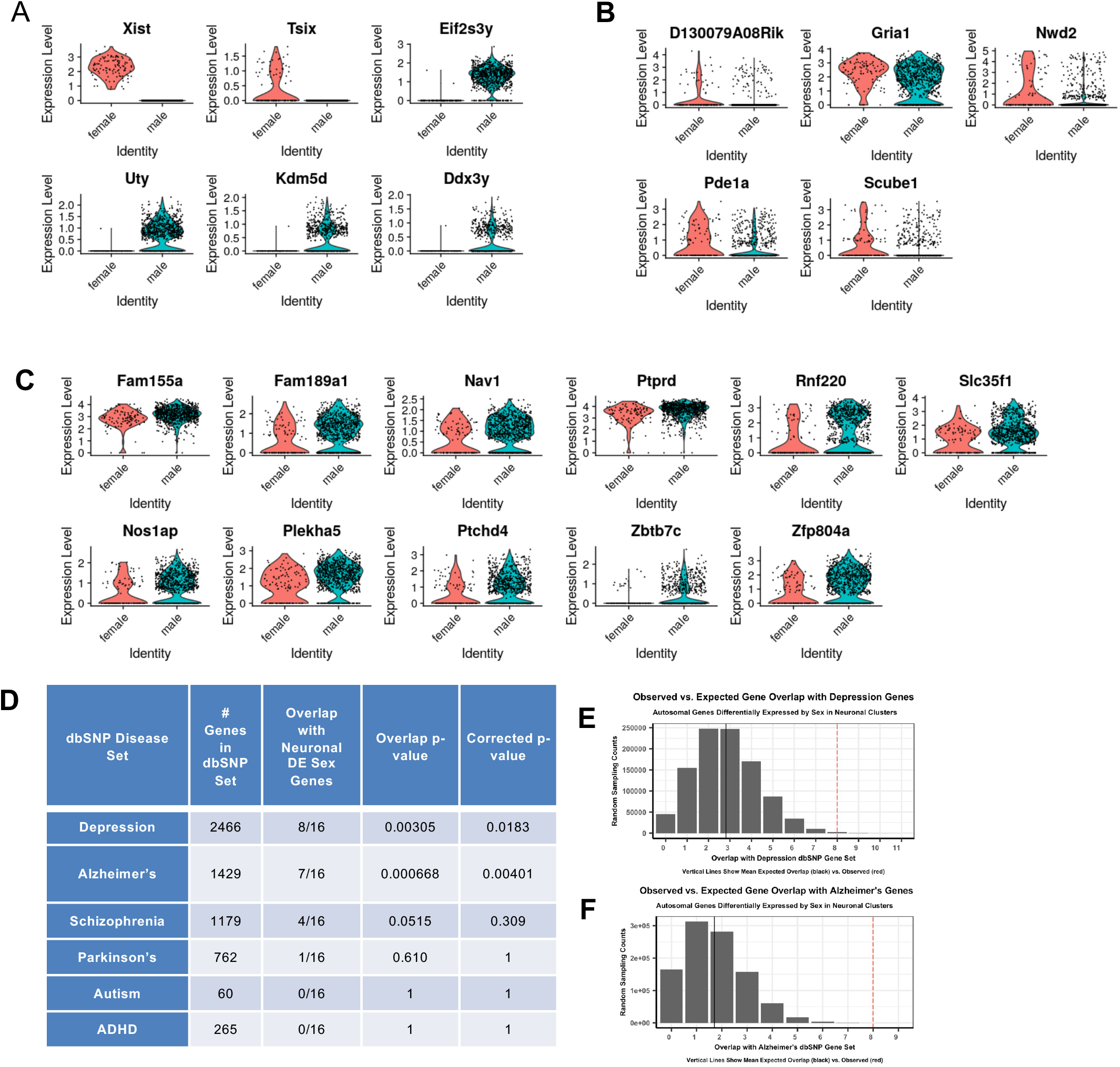
Sex bias in gene expression within Gad2+ populations from the LHb. **(A)** Expression of known female and male-specific genes within nuclei tagged by LMO barcodes added to nuclei isolated from female or male mice, respectively. **(B**,**C)** Differential expression of genes in Gad2+ LHb nuclei from female or male mice. Expression higher in female **(B)**, expression higher in male **(C). (D)** Table of comparison between sex-biased genes and dbSNP genes in the disease categories shown. The p-values were calculated as described in the results section and were Bonferroni corrected for multiple hypothesis testing. Observed versus expected overlap between sex-biased genes and dbSNP genes for depression **(E)** and Alzheimer’s **(F)**. Dotted red line, observed, black line, expected.

Excluding the 6 genes we used for sex classification, we found 17 additional genes that were significantly differentially expressed between *Gad2*+ neurons of male and female mice **(Table S5)**. One of these genes, *Gabra3*, is found on the X chromosome, and was expressed more highly in males (**Fig. S11A**). The other 16 are autosomal genes, five of which were expressed significantly more highly in Gad2+ nuclei from female mice (**Fig. 6B**) whereas the other eleven were expressed more highly in Gad2+ nuclei from male mice (**Fig. 6C**). We confirmed that the differences in gene expression did not arise from skewing of cell recovery across the five *Rbfox3+* clusters and that both male and female classified nuclei expressed similar levels of *Gad2* (**Fig. S11**).

The LHb has been implicated in stress-related disorders of the brain including major depressive disorder (MDD), and many of these disorders have a sex bias. We therefore asked whether the genes we identified as sex-differentially expression in Gad2+ neurons showed genetic association with psychiatric or neurological disorders. Using all genes with average expression above 0.25 from our LHb neuronal clusters as background, a one-sided Fisher’s exact test was used to determine if our sex-biased genes had significant enrichment within the dbSNP database of genes linked to a variety of neuronal phenotypes (Li et al., 2012). We computed the overlap of our 16 autosomal sex-biased genes with 6 categories of neuronal phenotypes from the dbSNP database, which we also filtered for autosomal genes, and calculated the statistical significance of this overlap by random sampling the LHb neuronal background set 1,000,000 times and applying Fisher’s Exact test between observed and expected overlaps. These analyses revealed significant overlap of LHb Gad2+ sex-biased genes with Depression and Alzheimer’s disease, which are both known to have female bias in the human population (**Fig. 6D-E)** (Laws et al., 2018).

### Ntng2 as a marker of Gad2/Slc17a6 double positive LHb neurons

To validate the transcriptomes from our snRNA-seq analyses and to identify a potential marker for this unique population of cells, we searched for uniquely expressed marker genes in the *Gad2*+ cells. We noticed that the gene for Netrin G2, *Ntng2*, appeared near the top of the DE gene lists for up-regulated genes when we compared these cells within our dataset of Gad2+ neurons (**Fig. 4A; Table S1**) whereas *Ntng1* was significantly down-regulated. The netrins are known to mediate axonal guidance and thus raised our interest in the possibility these molecules could define the projection targets of the *Gad2/Slc17a6*+ LHb neurons. *Ntng2* was also significantly upregulated when compared to other LHb neurons from the Hashikawa et al., 2020 dataset suggesting it could define these cells regionally (**Fig. 5D; Table S2**).

We performed in situ labeling using RNAScope FISH probes for *Gad2, Ntng1*, and *Ntng2* to determine the anatomical expression in LHb tissue and confirm the expression of *Ntng2* within *Gad2*+ neurons **(Fig. 7; Fig S12)**. We found distinct patterns of *Ntng1* and *Ntng2* expression in and around the habenular complex, with *Ntng1* expressed predominantly in the surrounding regions including the paraventricular nucleus of the thalamus (PVT) (**Fig. 7A-C, E)**. Interestingly, though *Ntng2* was densely expressed in the MHb, it showed a scattered distribution into the medial part of the LHb **(Fig. 7A-B, D, E)** where nearly all the *Gad2*+ neurons were observed to co-express *Ntng2* **(Fig. 7D, F)**. These findings validate the results from snRNA-seq study and provide a new marker for this cell type within the LHb.

**Figure 7:**
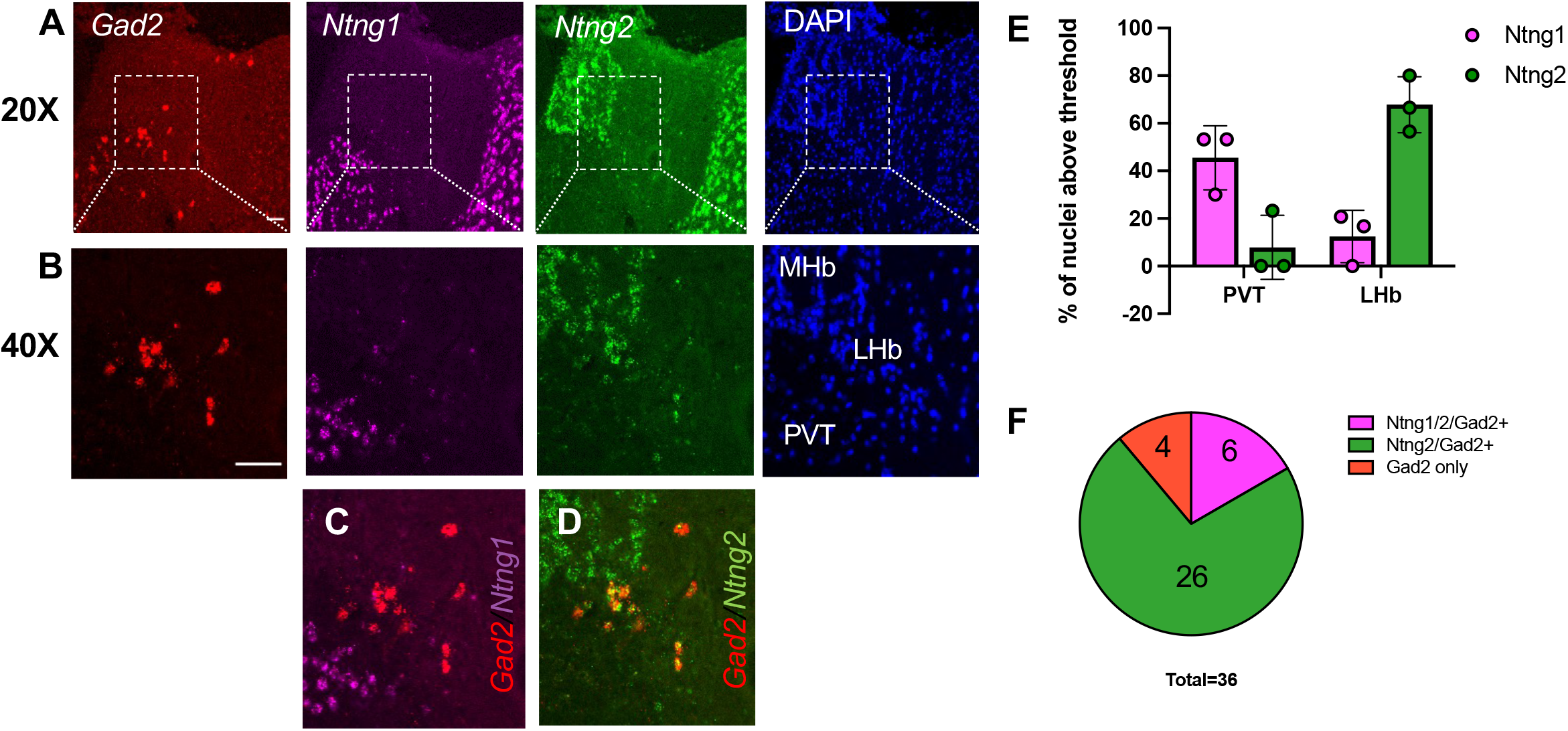
*Ntng2* is co-expressed in *Gad2*-positive neurons in the medial subnucleus of the LHb. **(A)** Representative RNAscope FISH images for *Gad2, Ntng1*, and *Ntng2* as well as DAPI at 20X The medial subnucleus of the LHb on each image is centered within the dotted white box, which is shown at 40X in **(B)**. The regions that correspond to the MHb, LHb, and PVT are labeled on the DAPI image. **(C)** Nonoverlap of *Gad2* with *Ntng1* and **(D)** overlap with *Ntng2* in the LHb. **(E)** Quantification of the percent of positive nuclei for each individual brain section from thresholding in **Fig. S12**. 30 cells/image/channel from 3 brains. **(F)** Percentage of the total of all *Gad2*+ positive nuclei from images of 3 brains that co-express *Ntng1, Ntng2*, or both. Scale bar: 40 μm.

## DISCUSSION

The widespread use of single cell sequencing methods has substantially advanced knowledge about the complexity of brain regions and variation within what used to be considered traditional cell types. However, rare cell types remain challenging to characterize in single cell analyses because they are found in too low abundance to drive subclusters with sufficient power to permit differential expression analysis. One strategy that has been successfully used to identify heterogeneity within rare cell populations such as GABAergic interneurons is to use genetic enrichment strategies prior to single cell sequencing (Gallegos et al., 2022; Munoz-Manchado et al., 2018). Here we used transgenic expression of the nuclear envelope protein Sun1-GFP to create a means to purify nuclei of *Gad2*+ neurons from the LHb for snRNA-seq. Consistent with the sparse pattern of *Gad2* reactivity on tissue sections, GFP+ nuclei comprised only ∼3% of all the FAC-sorted, DAPI+ nuclei in our microdissected samples. We recovered less than 700 of these cells from the LHb of each single mouse, which is consistent with *Gad2*+ cells comprising a few percent of the estimated 13,000 total LHb cells per mouse (Zhang & Oorschot, 2006). Given the low abundance of these cells, it is not surprising that they failed to be detected as a distinct cluster in either of two prior scRNA-seq datasets from mouse habenula (Hashikawa et al., 2020; Wallace et al., 2020). An scRNA-seq study of zebrafish habenula did detect a single *Gad2*+ cluster, however cell type variation within this cluster was not characterized (Pandey et al., 2018). This is important because in addition to the *Gad2+/Slc17a6+* cells we characterized here, there are *Gad2+/Slc17a6-*neurons both within and directly adjacent to the habenular complex that are known to have gene expression programs (*Pvalb+, Sst+*) and functions (local inhibition) that are similar to well characterized neocortical inhibitory interneurons (Webster et al., 2020; Yuste et al., 2020).

The enrichment we performed allowed us to sequence enough cells to compare different classes of *Gad2*+ cells among the isolated nuclei. Our analysis resolved 5 clusters of FANS enriched neurons with significantly differential programs of gene expression. *Gad2* mRNA was only weakly expressed within some of these clusters (clusters 1 and 5; **Fig. S5**), however this was presumably sufficient to drive enough Sun1-GFP expression from the Cre-dependent transgene to allow these cells to be sorted by FANS (**Fig. S1C**). When we compared expression of the top differential genes in each cluster with *in situ* localization patterns from the Allen Brain Atlas from the Allen Institute only cluster 2 markers strongly overlapped the distribution of the *Gad2+/Slc17a6+* population in the medial subnucleus of the LHb (**Fig. 1F, Fig. S1B**). Clusters 1 and 3 appear to be derived primarily from *Gad2*-driven Sun1-GFP transgene expression (**Fig. S1A**) in the dorsal thalamic regions immediately flanking the LHb, which likely contaminated our microdissection. Cluster 5 contains marker genes that are widely expressed in MHb as well as in neurons scattered through LHb. Given that the MHb shows no detectable *Gad2* expression (**Fig. 1E**) or *Gad2*-driven Sun1-GFP transgene expression (**Fig S1A**), cluster 5 is likely to come from cells in the LHb that share some similarities in gene expression with MHb neurons. Finally, cluster 4’s expression of canonical GABAergic genes including *Pvalb, Slc32a1*, and *Gad1*, as well as *Gad2*, suggest that this is a local inhibitory interneuron population. There are *Pvalb*+ neurons within the LHb (**Fig. 1F**) however parvalbumin only colocalizes with GABA in a small percentage of LHb *Pvalb*+ neurons, and these are localized in lateral but not the medial regions of the LHb (Webster et al., 2020). Thus, if these cells did come from the habenula, the Cluster 4 neurons we isolated and sequenced are likely to represent this lateral LHb *Gad2*+ population, where the inhibitory function of *Pvalb*+ neurons has been previously characterized (Webster et al., 2020).

Neurons that co-express markers of more than one neurotransmitter system have now been found in many regions of the brain, in sharp contrast to the historical “one neuron, one transmitter” principle (Root et al., 2018). However, the cellular and molecular biology of dual transmitter release remains to be fully understood. *Gad2*+ neurons in the medial subnucleus of the LHb have been shown to mediate both local inhibitory neurotransmission in the LHb (Flanigan et al., 2020) and long distance excitatory neurotransmission (Quina et al., 2020).

Importantly, among our purified *Gad2*+ LHb neurons, the cells in cluster 2 were unique for their expression of *Hcrtr2*, the gene encoding the orexin receptor ORXR2, which further validates these neurons as the same cells determined by Flanigan et al. (Flanigan et al., 2020) to be locally inhibitory within the LHb. We observed both by single cell sequencing (**Fig. S5**) analysis of cluster 2 and by *in situ* hybridization (**Fig. 3)** that *Gad2+* in the medial subnucleus of the LHb co-express the vesicular glutamatergic transporter *Slc17a6*, encoding VGLUT2. We confirmed previous reports that these cells fail to co-express the vesicular GABA transporter VGAT, encoded by *Slc32a1* (Quina et al., 2020; Wallace et al., 2020), but we did find colocalization of *Gad2*/+*Slc17a6+* cells in LHb with *Slc6a1*, encoding the plasma membrane GABA transporter (**Fig. 3)**. One possibility is that reversal of this plasma membrane GABA transporter could be used for non-vesicular GABA release, explaining how these neurons can drive inhibition (Richerson & Wu, 2003). Alternatively, other transporters could package GABA in vesicles; for example, midbrain dopamine neurons use as *Slc18a2*, encoding VMAT2, to package GABA into synaptic vesicles for release (Tritsch et al., 2012). When mechanisms are discovered that can be used by neurons to release multiple transmitters, the transcriptome we have characterized for the *Gad2+/Slc17a6+* cells in cluster 2 will be a useful resource to better understand the physiological functions of these cells in LHb-coupled circuits.

One of the most powerful applications of scRNA-seq is linking molecular signatures of distinct cell types with the physiology assigned to a given brain region (Armand et al., 2021). Here we demonstrated that we could use alignment of identifiable cell types in clusters to integrate our dataset enriched for rare *Gad2+* cells of the LHb into a prior dataset containing unbiased representation of abundant LHb cell types (Hashikawa et al., 2020). This allowed us to find LHb genes that are preferentially expressed in the *Gad2+* population, suggesting LHb functions that map to these neurons. For example, the LHb is a hub that integrates upstream activity from limbic forebrain and basal ganglia and controls the downstream activity of midbrain neuromodulatory nuclei (Hu et al., 2020). Though many studies have focused on the importance of LHb projections in the control of dopamine systems in the brain, the *Gad2+/Slc17a6+* neurons are thought to project primarily to serotonergic neurons in the dorsal and median raphe nuclei (DR, MnR) (Quina et al., 2020). We find that all the *Gad2*+ neurons we isolated not only express multiple serotonin receptors (**Fig. 4E**) but also express them at higher levels compared with other types of LHb neurons (**Fig. S9**). The LHb receives a dense serotonergic projection back from the DR (Metzger et al., 2021), suggesting that these neurons may be engaged in a bidirectional feedback loop that contributes to serotonergic regulation of the LHb’s role in pain perception, addiction, approach/avoidance behaviors, and motivation (Tchenio et al., 2016). Further consistent with roles for *Gad2*+ neurons of the LHb in psychiatric disorders, we found significant overrepresentation of a number of genes associated with depression (Howard et al., 2019) in the *Gad2+/Slc17a6+* population compared with other LHb neurons (**Fig. 5F**), and genes that showed sex-differential expression in LHb *Gad2+* cells were more likely than chance to be associated with depression and Alzheimer’s risk genes (**Fig. 6D,E**). Future studies that use intersectional strategies such as the unique colocalization we detected between *Gad2* and *Ntng2* to target this population for transgene expression will allow refined circuit studies that directly test the function of this cell population in the LHb.

## METHODS

### Animals

We used adult (>P60) male and female mice, and all experiments were conducted in accordance with an animal protocol approved by the Duke University Institutional Animal Care and Use Committee. We used CD1(ICR) (RRID:IMSR_CRL:022) and C57BL6/J mice (RRID:IMSR_JAX:000664) for in situ hybridization. For nuclear isolation, we crossed homozygous *Gad2*-IRES-Cre mice (*Gad2*^*tm2(cre)Zjh*^/J, RRID: IMSR_JAX:010802) (Taniguchi et al., 2011) with homozygous mice expressing a Cre-inducible Sun1-myc-sfGFP transgene, also known as INTACT (B6;129-*Gt(ROSA)26Sor*^*tm5(CAG-Sun1/sfGFP)Nat*^/J, RRID: IMSR_JAX: -21039) (Mo et al., 2015) to generate dual *Gad2*-Cre/INTACT heterozygotes (HET) for nuclear isolation.

### Nuclear isolation of Gad2+ nuclei from LHb

Fresh LHb was dissected by punch biopsy from 1mm sections bilaterally from *Gad2*-Cre/INTACT HET mice (n=7 male, n=3 female). Once the tissue was isolated, LHb samples were pooled separately by sex, and tissue was dounce homogenized prior to nuclear isolation on an Optiprep gradient as previously described (Gallegos et al., 2022). Nuclei were then re-suspended, washed in homogenization buffer, and incubated with MULTI-seq lipid-modified oligos (LMOs) (McGinnis et al., 2019) to barcode nuclei from either male (barcode 1) or female (barcode 2) mice respectively for retrospective analysis of sex-specific differences in gene expression. LMOs were added at a ratio of 10:1 oligo barcode to molecular (lipid) anchor and DAPI. The male and female LHb nuclei were then pooled and sequenced together to overcome potential batch effects during 10X sequencing.

### Fluorescence activated nuclear sorting and single nucleus RNA-sequencing (FANS-snRNA-seq)

*Gad2*+ nuclei were isolated using Fluorescent-Activated Nuclear Sorting (FANS) gating on double positivity for DAPI and GFP and sorted into homogenization buffer. Flow cytometry was performed in the Duke Human Vaccine Institute research flow cytometry shared resource facility. Purified nuclei were run on a Beckton Dickinson FACSAria IIu 70μm nozzle with a sheath pressure of 70psi. Nuclei were separated from debris by gating a histogram of DAPI fluorescence around the singlet peak. GFP positive nuclei were defined and gated from events exceeding the minimum density point between the center of the GFP- and GFP+ populations. Out of 242,8333 DAPI-positive events, 3% (7285) were double DAPI/GFP+ nuclei that were sorted into a single well of a 96-well-plate for 10X Genomics snRNA-Seq. 10X Genomics 3’ Gene Expression (v3 chemistry) droplet generation and library construction were performed at the Duke DHVI Sequencing Facility and sequenced on Illumina Nextseq 550 in mid output mode. Data are deposited at GEO datasets at GSE179198.

### Data analysis for snRNA-seq

Raw BCL files were converted to fastqs using CellRanger v3.0.2 mkfastq. Fastq files were then aligned to the mm10-3.0.0_pre-mRNA reference transcriptome and a count matrix was generated using CellRanger v3.0.2 count. This count matrix was used as the input to Seurat V417 (Johnson et al., 2021) for downstream analysis. We identified the top 2000 highly variable features to use for principal component analysis (PCA) and the top 10 principal components were used to perform cell clustering at a resolution of 0.5 with the non-linear dimensional reduction technique, Uniform Manifold Approximation and Projection (UMAP). Filtered feature matrix files were imported into Seurat V417. Single cells were filtered with a minimum of 200 features and data was normalized for each cell by the total expression multiplied by a scale factor of 10,000 and a log-transformation of the result. As in a previous study of LHb (Hashikawa et al., 2020), we used the following marker genes to identify cell types: *Tac2* for medial habenula (MHb), *Pcdh10* for LHb, *Rbfox3* for neurons, *Tmem119, Cx3cr1*, and *Pros1* for microglia, *Cldn5* and *Flt1* for endothelial cells, *Tagln* for mural cells, *Olig1* and *Mog* for oligodendrocytes, *Aldoc* and *Slc6a11* for astrocytes, and *Pdgfra* and *Gpr117* for oligodendrocyte precursor cells. We filtered for the neuronal clusters (clusters 1-5) with high abundance of *Rbfox3* transcripts (n=1491 nuclei, 3416.6 avg genes per nucleus, 8201.9 avg counts per nucleus). Differential expression was performed between clusters using a Wilcoxon rank Sums test using default settings in Seurat of log fold change greater than 0.25 and p-values less than 0.05. Gene ontology (GO) analysis was performed using Enrichr (Chen et al., 2013). Differentially expressed (DE) genes were analyzed for their significance within top GO terms categorized by cellular function (CC), molecular function (MF), and biological process (BP) terms using Fisher’s exact significance test.

### RNAscope in situ hybridization

Mice were deeply anesthetized with isoflurane, and brains were harvested and flash-frozen in isopentane/dry ice baths. 20μM coronal slices containing the LHb were sectioned on a cryostat and mounted on Super Frost Plus slides. We performed RNAscope fluorescent *in situ* hybridization as previously described (Wang et al., 2018) using the following Biotechne ACD probes: *Gad2* (Cat.: 439371-C3), *Slc6a1* (Cat.: 444071-C2), *Slc17a6* (Cat.: 319171), *Ntng1* (Cat.: 488871-C2), and *Ntng2* (Cat.: 585811). Sections were incubated with DAPI for 3 minutes, covered with ProLong Gold antifade and covered with coverslips. Z-stack images (10μM steps of 10 stacks) were taken with a Zeiss 880 inverted confocal microscope and visualized and analyzed in FIJI/ImageJ.

### Quantification of RNAscope images

Max projection images at 40X centered on the medial subnucleus of the LHb were obtained from three independent mice for each analysis, and quantification of signal intensity was performed in FIJI. To establish the background signal in each channel on each slide, an investigator used the DAPI channel to randomly choose 15 equal sized circular ROIs in cell-poor regions of the image. These ROIs were lifted to each of the other channels in the image stack, and the background signal intensity in each channel was measured in this common ROI set. To quantify multiple *in situ* signals in single cells, the investigator then used the DAPI image to randomly choose either 40 (for **Fig. 3**) or 30 (for **Fig. 7**) equal sized ROIs overlapping nuclei in any given brain region (MHb, LHb, or PVT). Again, the intensity of *in situ* signal was measured for this common ROI set in each channel from the stack. Finally, because *Gad2*+ neurons are rare and not all were captured in the random assessment, to specifically determine the overlap of *Gad2* with other genes of interest, an investigator first identified ROIs overlapping all *Gad2*+ (positive) cells on a slide and then lifted this common ROI set to measure the intensity of all other channels in the same image stack. In all cases, signal intensity in each ROI was divided by the average of the background signal for that channel on that slide to give a normalized intensity with the background scaled to 1. We defined the average background signal plus 2 standard deviations as the threshold to call an *in situ* signal positive.

### Integration with scRNA-seq data of other LHb cell types

A total LHb scRNA-seq dataset for comparison to our data was taken from Hashikawa et al., 2020 (Hashikawa et al., 2020). These data were obtained from GEO accession number GSE137478 and processed in Seurat according to our previous pipeline. 40 PCs were used to perform clustering and further analysis of the integrated dataset. Labels of cells from our independent dataset were used to track cells through integration. Differential expression was conducted as above, but because of the large number of sequenced cells in this dataset, we used a non-parametric Wilcoxon Rank Sum test with an adjusted p-value ≤ 10e^-32^.

### Sex-specific gene analysis

For sex-specific analysis, original MULTI-seq LMO calls from male or female mice were read in as a .csv and added to the raw count Seurat Object as a metadata variable in a barcode-specific manner. For nuclei in the *Rbfox3+* neuronal clusters, LMO calls (LMO1/Male n=391, LMO2/Female n= 135) were converted into cell Identities using the Idents command and differential expression was performed using a Wilcoxon Rank Sum test via the FindMarkers command as above. We confirmed sex-specific enrichment of female-specific X chromosome transcripts (*Xist, Tsix*) and male-specific Y chromosome transcripts (*Eif2s3y, Uty, Kdm5d, Ddx3y*) by female and male LMO barcodes respectively. To account for dropout of any single gene, we used the relative abundance of these transcripts to create an expression-based classifying for identifying sex. Nuclei were classified as male or female if they passed a binned kernel density estimate threshold of 20 for LMO barcode reads or had sex-chromosome gene expression signatures above a computed confidence score of 0.66 (with greater than 2 UMI reads across the sex-specific gene set). After these classifications, 107 female and 816 male nuclei were confidently identified, and these expanded calls were integrated as a metadata variable and used to create cell identities for differential expression in Seurat as above.

### Sex-specific Genes Cross-comparison to Neuronal Phenotype GWAS Gene Sets

Using all genes in our data set with average expression above 0.25 from our LHb neuronal clusters as background, a one-sided Fisher’s exact test with Bonferroni correction was used to determine if our sex differentially-expressed genes had significant enrichment within the dbSNP database of genes linked to a variety of neuronal phenotypes (Li et al., 2012) We computed the overlap of our 16 autosomal sex biased genes with 6 categories of neuronal phenotypes from the dbSNP database, which we also filtered for autosomal genes, and calculated the statistical significance of this overlap by random-sampling the LHb neuronal background set 1,000,000 times and applying Fisher’s Exact test between observed and expected overlaps. These analyses revealed significant overlap of LHb sex-biased genes with Depression and Alzheimer’s disease, and non-significant overlap with Schizophrenia, Parkinson’s, ADHD, and Autism.

## Acknowledgements

We thank Mariah Hazlett for critical reading of the manuscript and Melyssa Minto for support with the bioinformatics. This work was supported by NIH grant R01DA047115 (A.E.W.).

## SUPPLEMENTAL INFORMATION

**Figure S1:**
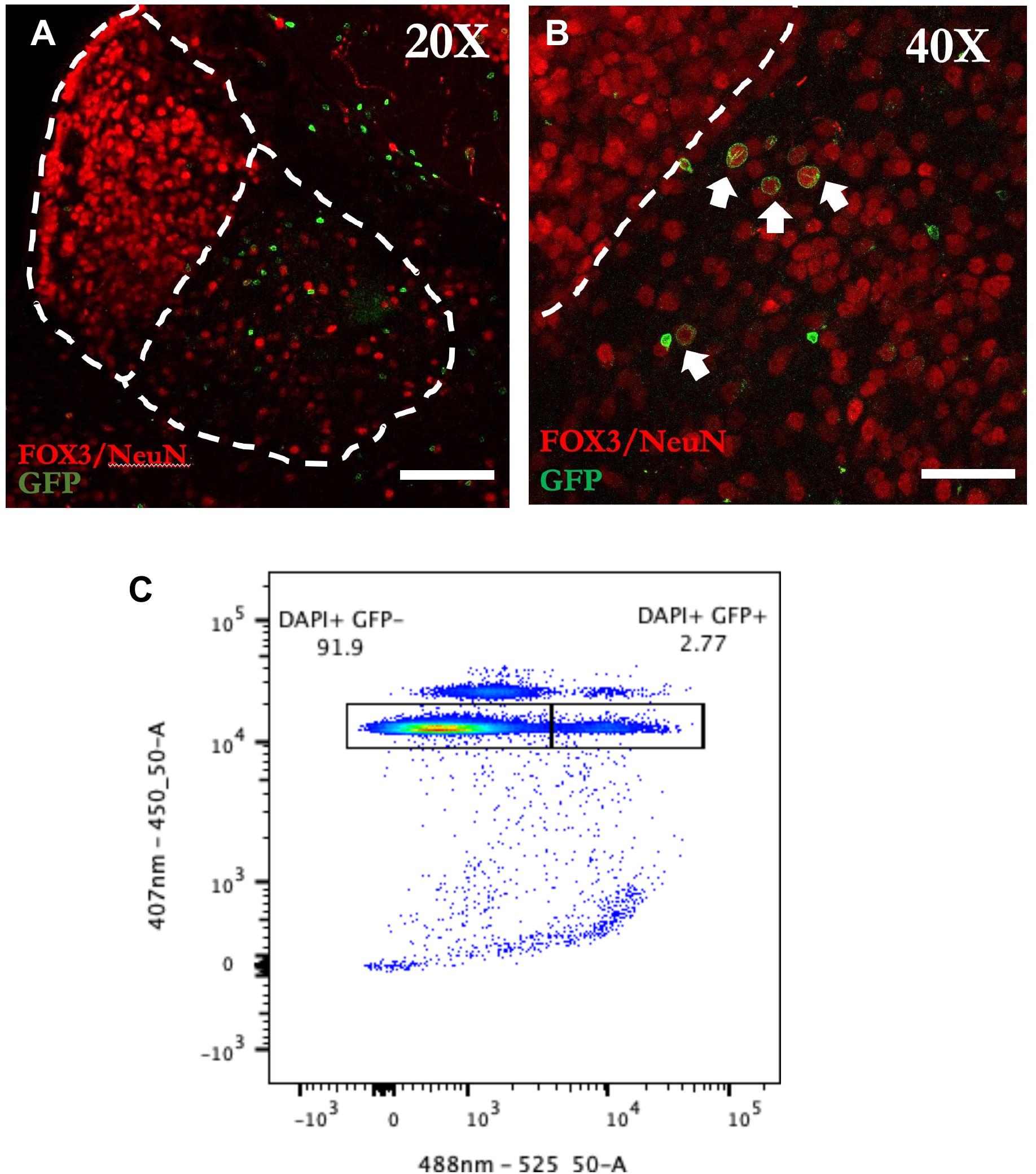
Genetic labeling and isolation of GAD2+ neurons from the LHb. (**A**,**B**) Sun1-GFP labeling around the nuclei of neurons in the medial subnucleus of the LHb in *Gad2*-Cre/INTACT HET mice (white arrows in **B**). Neuronal nuclei are labeled with the NeuN antibody, which recognizes the nuclear protein FOX3. Dotted white lines outline the medial (left side) and lateral (right side) habenula. Scale bar 200μm (**A**), 100μm **(B)**. (**C**) Representative FACS image showing gating parameters for DAPI+ single nuclei that are GFP- (left side) or GFP+ (right side). Values show percent of all events that are in the gating boxes. DAPI+ GFP+ nuclei were collected for single nucleus sequencing.

**Figure S2:**
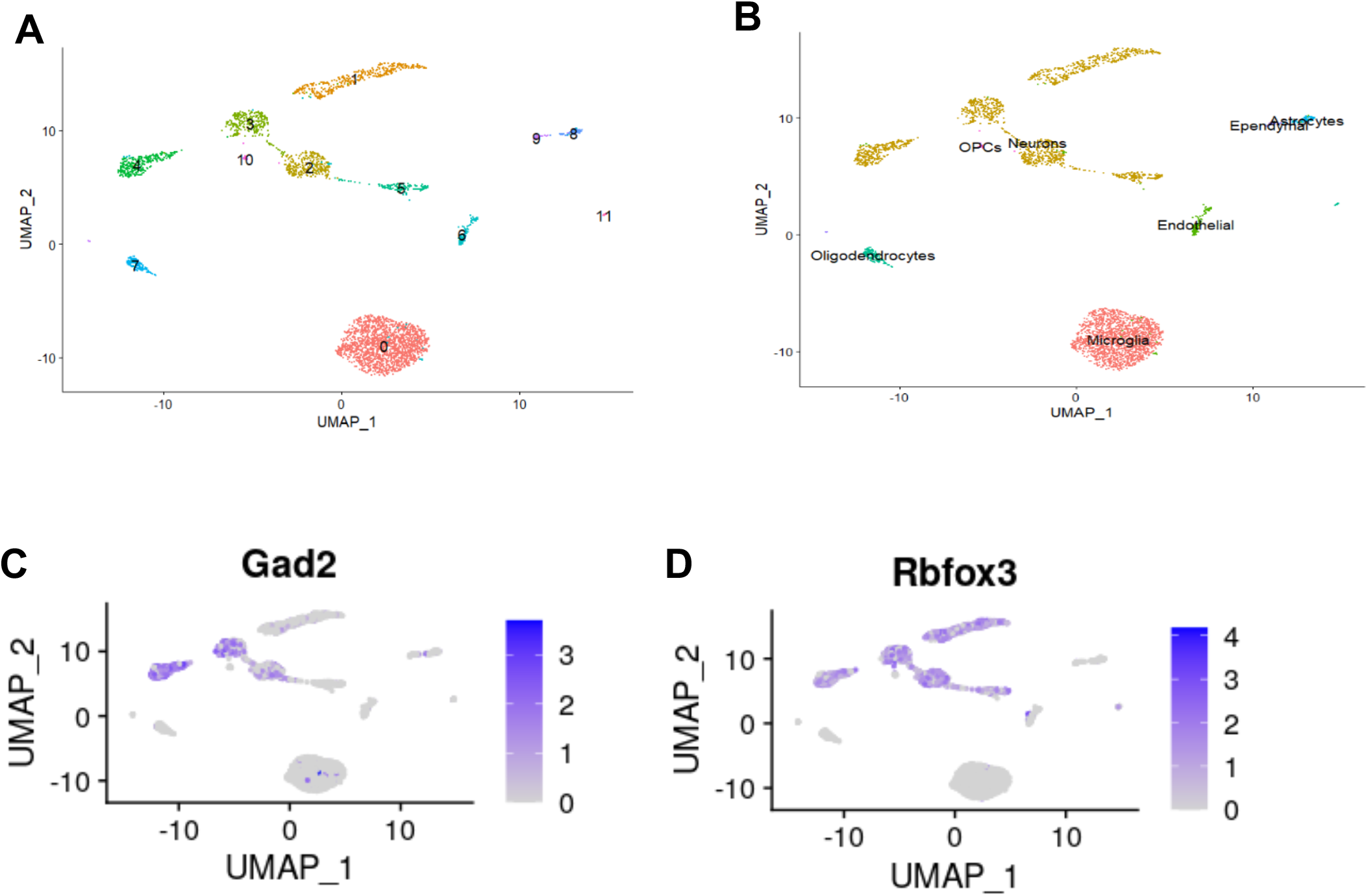
Identification of neuronal GAD2+ clusters. **(A)** UMAP clustering of GFP+ cells by FACS. **(B)** Color coding of cell types in each cluster based on marker genes (see methods). **(C**,**D)** Distribution of *Gad2* and the neuronal marker *Rbfox3* encoding FOX3 (see Fig **S1A,B**) within the 11 UMAP clusters.

**Figure S3:**
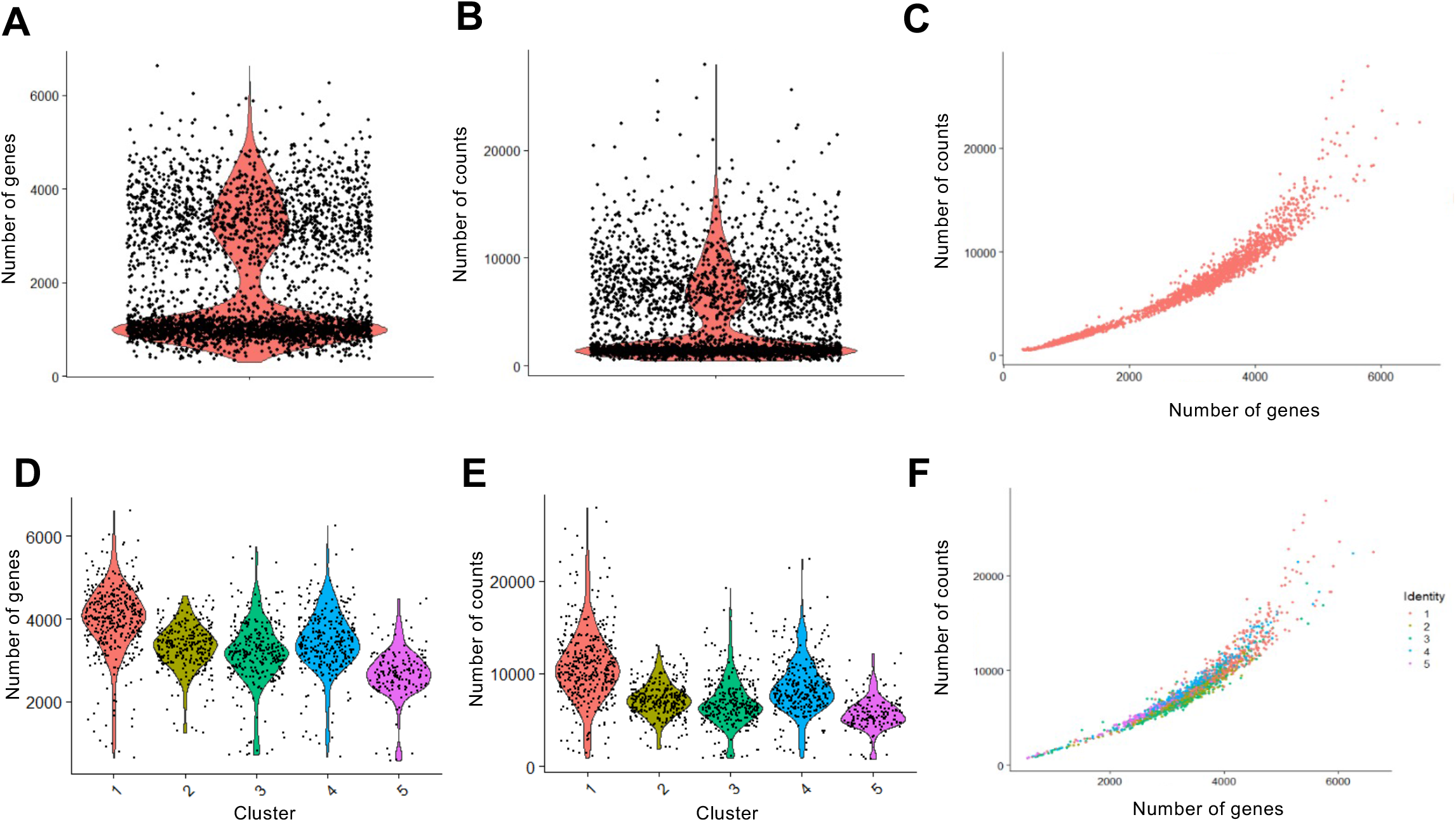
Similar QC features across the five neuronal (*Rbfox3*+) clusters. Violin plots showing the distribution of number of genes **(A)** and counts **(B)** per cell for entire dataset. **(C)** Feature scatter plot comparing the numbers of counts and genes per cell for the entire dataset. Violin plots showing the distribution of number of genes **(D)** and counts **(E)** per cell for the subsetted neuronal clusters 1-5. **(F)** Feature scatter plots comparing the numbers of counts and genes per cell for the subsetted neuronal cluster 1-5.

**Figure S4:**
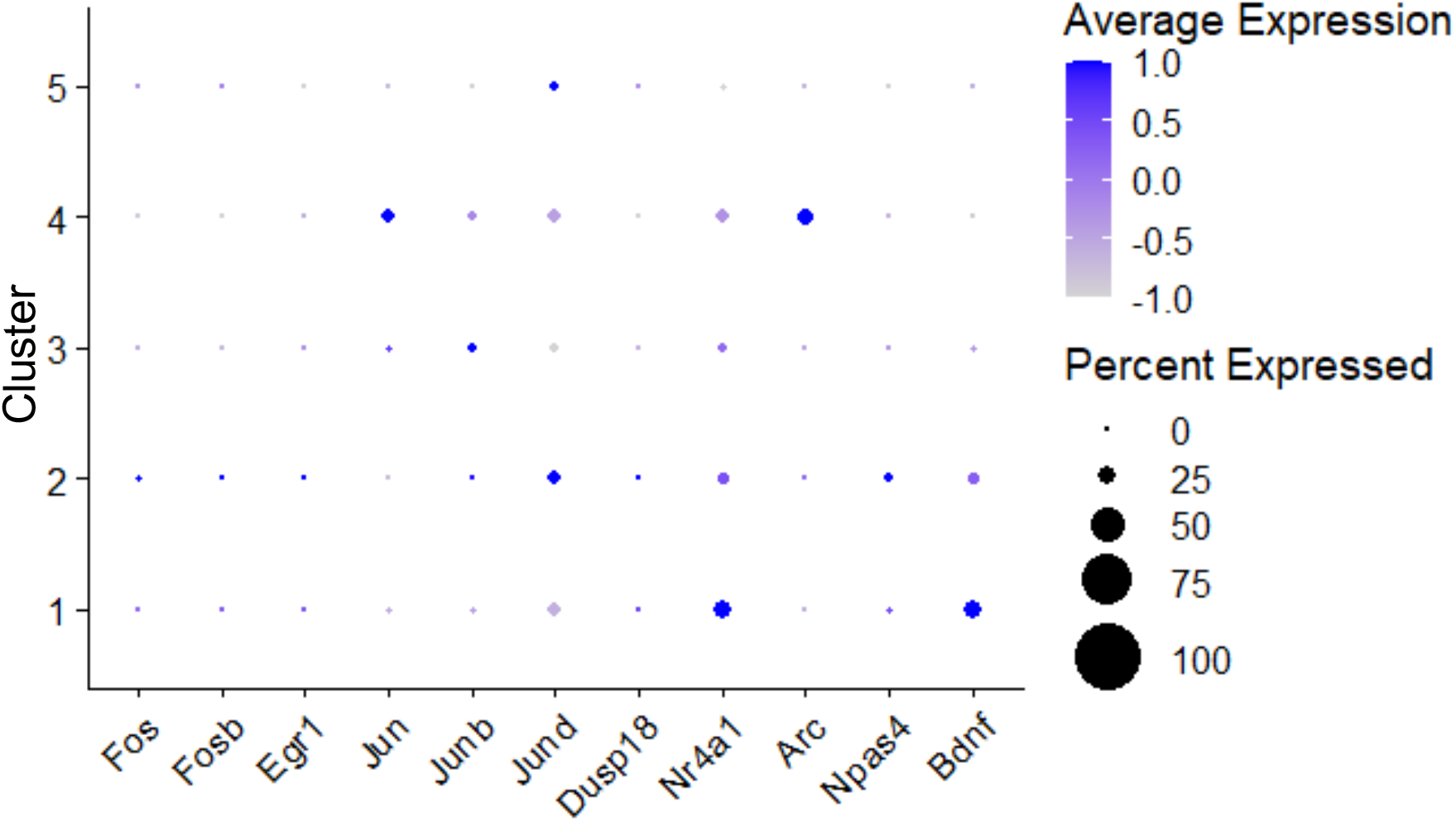
Activity-regulated genes are similarly detected in each of the five neuronal clusters. Dot plot shows scaled expression of the indicated activity-regulated genes in neuronal clusters 1-5.

**Figure S5:**
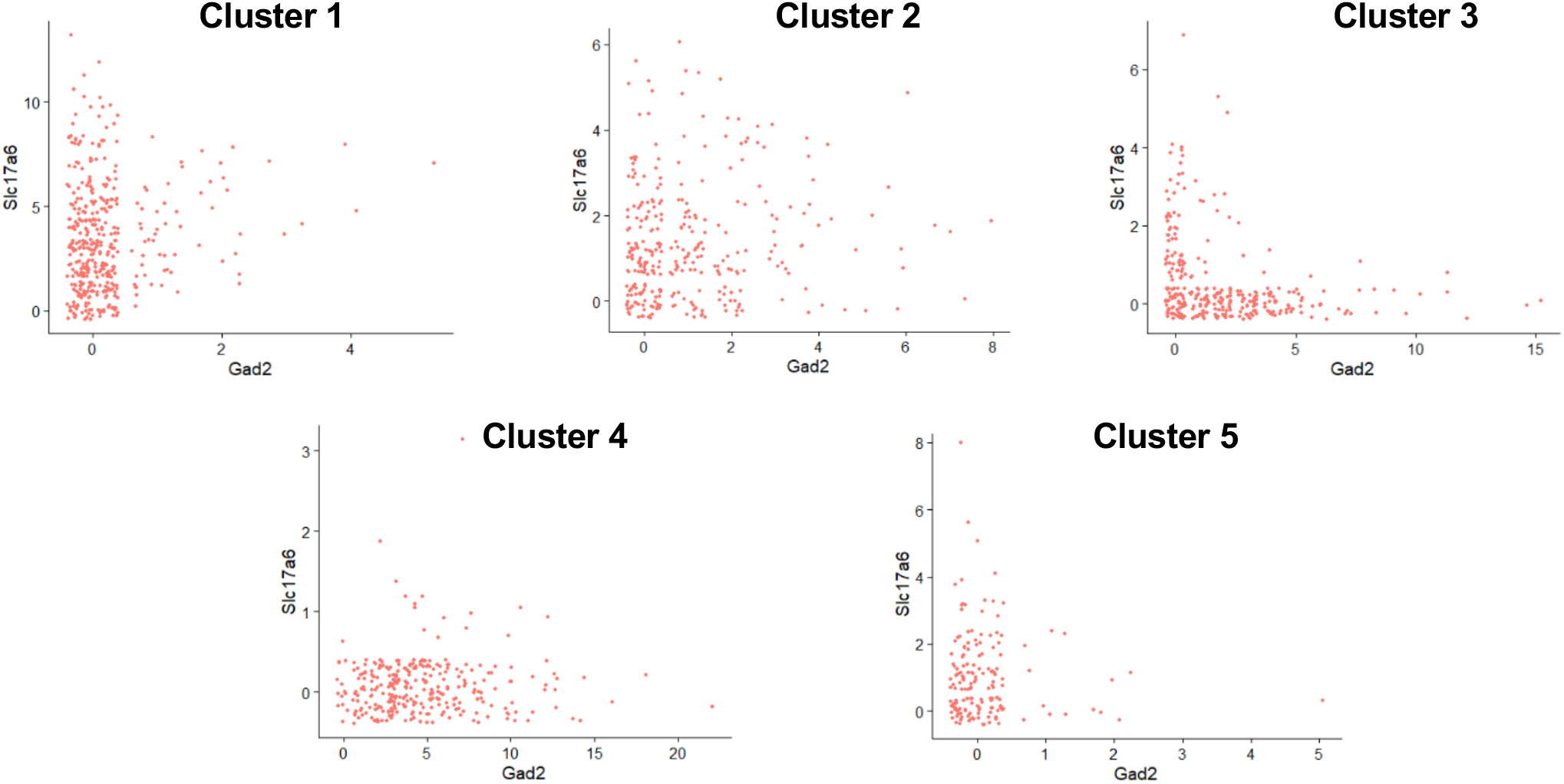
Cluster 2 shows strongest single cell co-expression of *Gad2* and *Slc17a6*. Feature scatter plots showing the expression levels of *Gad2* (x-axis) and *Slc17a6* (y-axis) in each single nucleus from snRNA-seq neuronal clusters 1-5.

**Figure S6:**
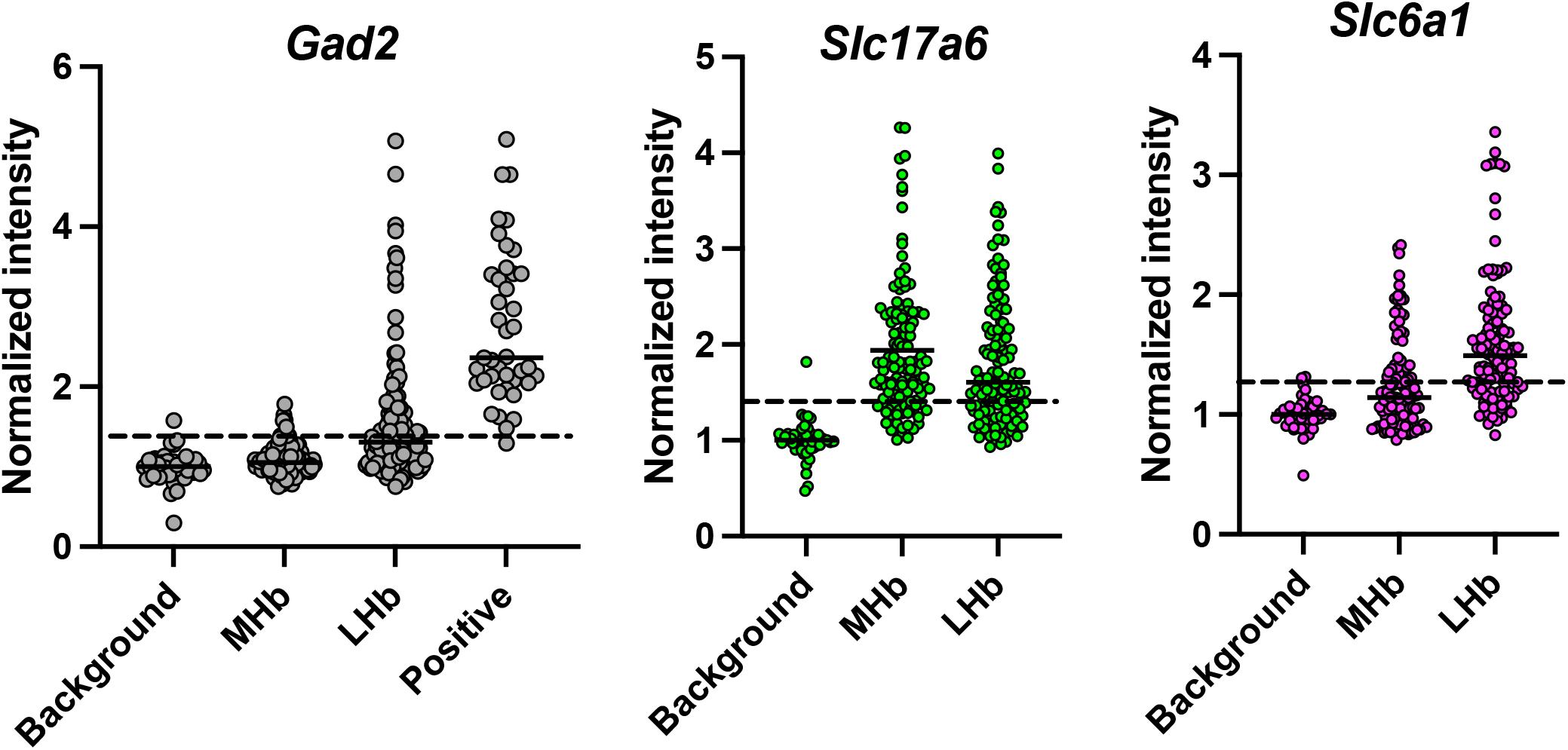
Threshold analysis of individual nuclear ROIs for each *in situ* channel in **Fig. 3**. Background shows 15 randomly chosen ROIs over cell-poor regions. The average background signal for each channel on each slide was used to normalize signal from all ROIs in the same stack. The dotted line is the average background signal + 2 standard deviations, which we defined as the threshold for calling a cell positive. Note that a small number of “background” ROIs are above the threshold because average plus 2SD accounts for 95% of the data within that set. We decided on this threshold by assessing the signal in ROIs over cells called “positive” by an investigator looking at the *Gad2* channel. All but one of these cells is above the threshold, suggesting the average+2SD threshold limits both false positives and false negatives in the quantification. Each circle is a single ROI. 15 background ROIs and 40 cells in each brain region were assessed for each of 3 images from independent brains. 52 Gad2-positive nuclei were assessed.

**Figure S7:**
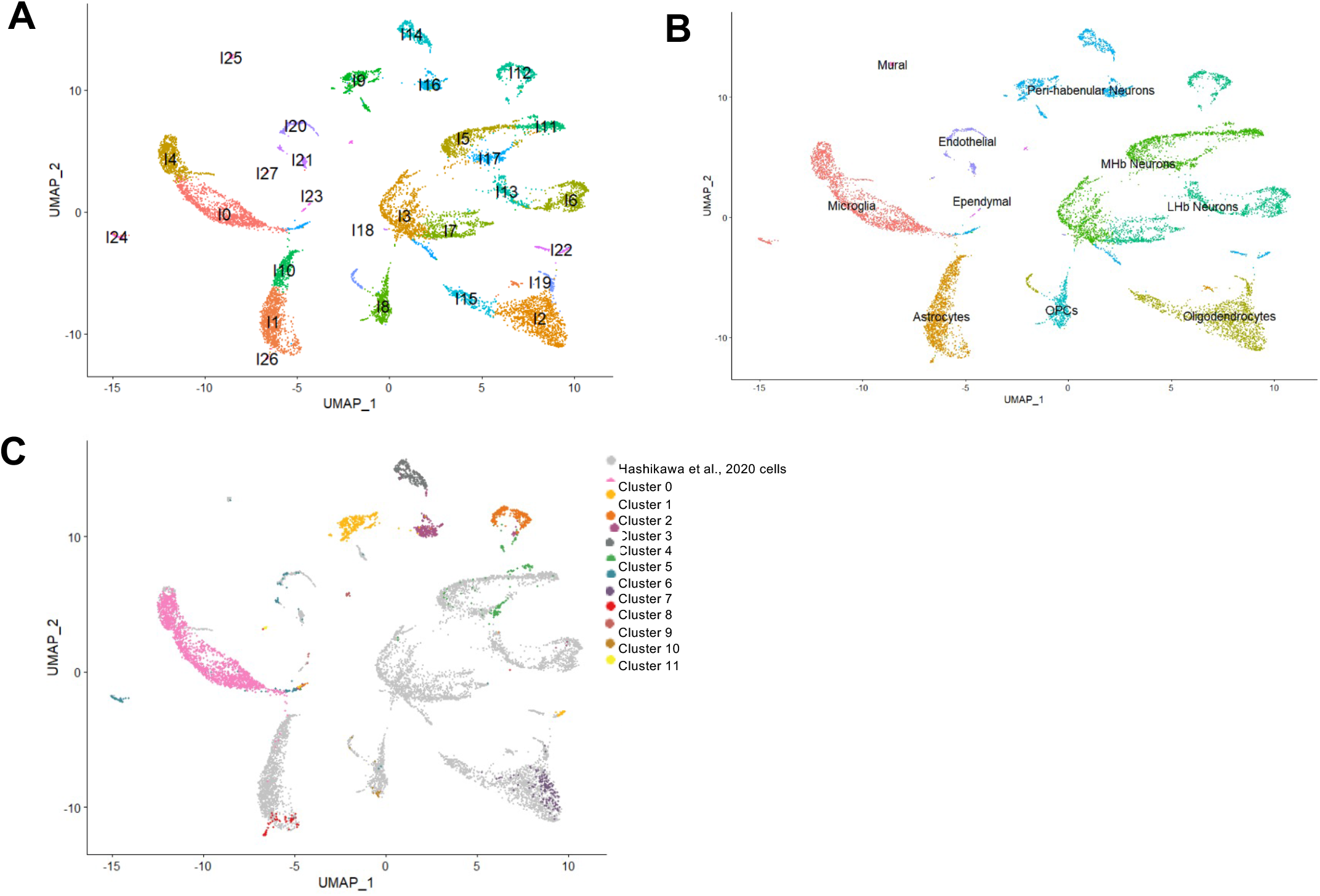
UMAP clustering of all LHb cells after integration of our Gad2+ snRNA-seq data with total habenula scRNA-seq data taken from Hashikawa et al., 2020 (3). Data shown by cluster number **(A)** and by cell type **(B). (C)** UMAP clustering highlighting the individual cells from our dataset (colored by original cluster number from **Fig. S2A**) versus the dataset in Hashikawa et al., 2020 (grey).

**Figure S8:**
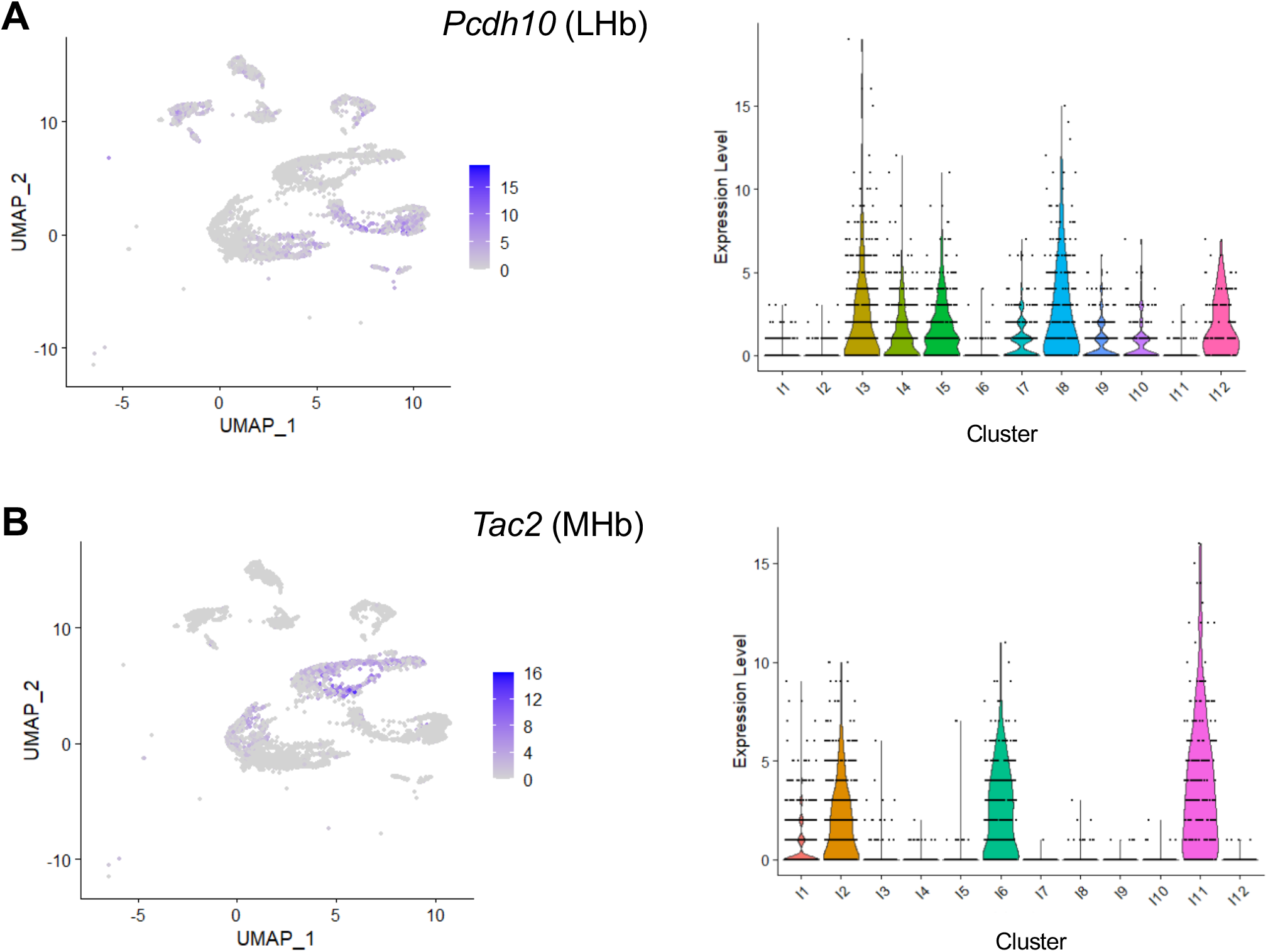
Marker identification of LHb and MHb neurons. **(A)** Feature plot showing the scaled expression of the LHb marker *Pcdh10* across all the clusters. Violin plot displaying raw counts per cell of *Pcdh10* expression for each cluster. **(B)** Feature plot showing the scaled expression of the MHb marker *Tac2* across all the clusters. Violin plot displaying raw counts per cell of *Tac2* expression for each cluster.

**Figure S9:**
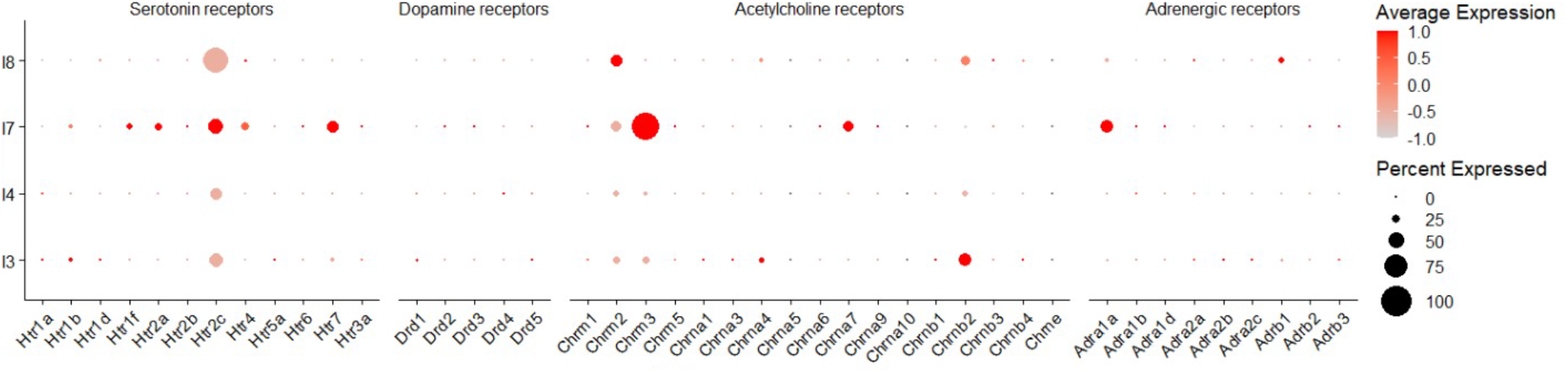
Dot plot showing the expression of genes for serotonin, dopamine, acetylcholine, and adrenergic receptors in clusters I3, I4, I7, and I8 from Figure 4A.

**Figure S10:**
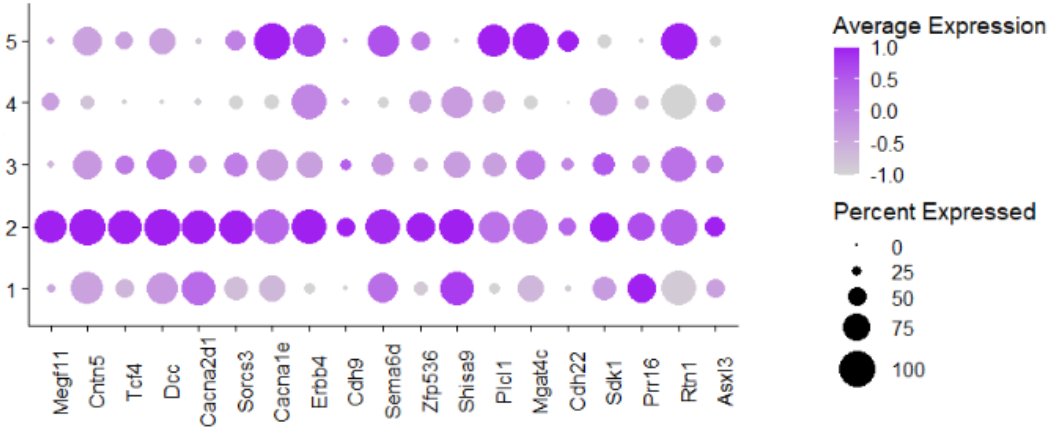
Dot plot showing scaled expression of MDD associated-genes that were preferentially expressed in cluster 2 (from Fig. 1) compared to all other neuronal clusters within our Gad2+ dataset.

**Figure S11:**
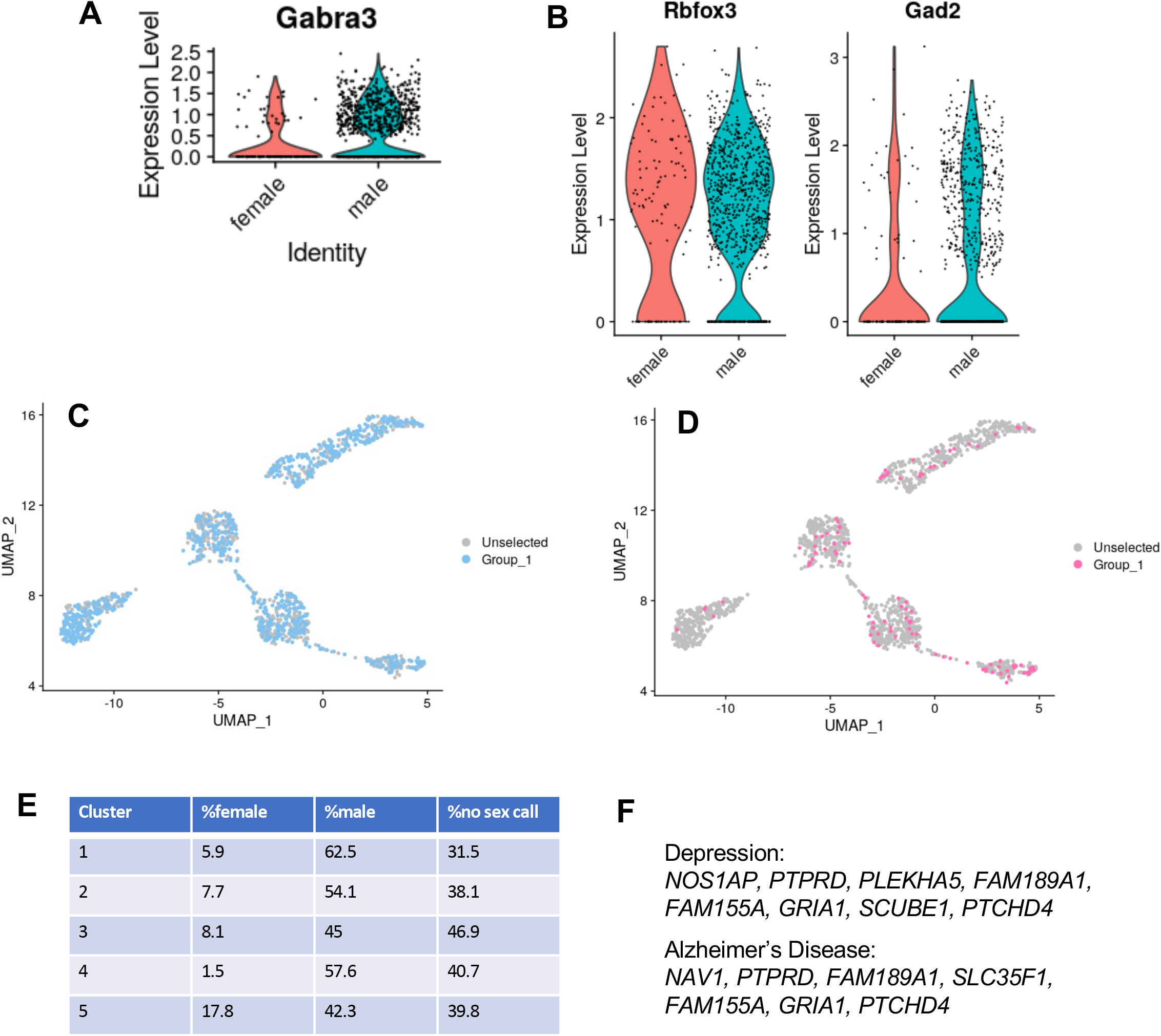
Gene expression analysis in nuclei recovered from male and female mice. **A)** Expression of the X-chromosome gene *Gabra3* in nuclei from female (red) and male (blue) mice. **B)** Similar expression of *Rbfox3* and *Gad2* in nuclei from male and female mice. **C, D) D**istribution of male (blue, **C**) and female (pink, **D**) neurons across the five *Gad2*+/*Rbfox3*+ clusters. **E)** Distribution of identified male and female nuclei, as well as nuclei that could not be sex categorized, across the five clusters. Of the genes that show higher expression in nuclei from female mice, only *Nwd2* and *D130079A08Rik* are selectively expressed in cluster 5 (see **Table S1**) raising the possibility that these two genes may appear sex-biased because of the skewed recovery of nuclei from females in this cluster. **F)** Sex-biased genes in Gad2+ neurons that are associated with depression and Alzheimer’s disease in dbSNP (**Fig. 6D**). Of the male-biased genes, only *Nos1ap1* has a cluster bias, and it is enriched in clusters 2,4, and 5 **(Table S1**). Of the female-biased genes, only *Gria1* has a cluster bias, and it is enriched in clusters 2,3, and 5.

**Figure S12:**
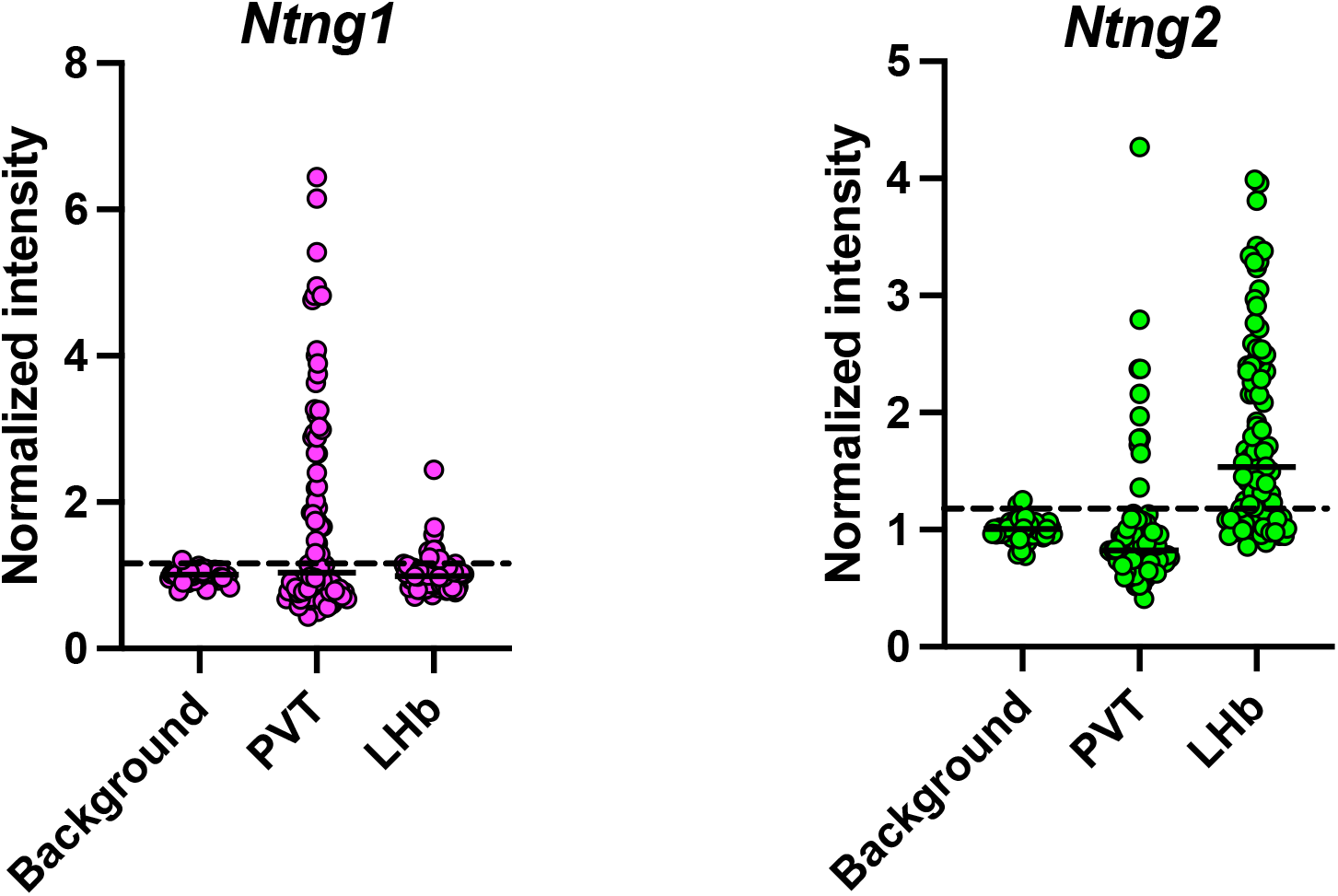
Threshold analysis of individual nuclear ROIs for each *in situ* channel in **Fig. 7**. Background shows 15 randomly chosen ROIs over cell-poor regions. The average background signal for each channel on each slide was used to normalize signal from all ROIs in the same stack. The dotted line is the average background signal + 2 standard deviations, which we defined as the threshold for calling a cell positive. Note that a small number of “background” ROIs are above the threshold because average plus 2SD accounts for 95% of the data within that set. Each circle is a single ROI. 15 background ROIs and 30 cells in each brain region were assessed for each of 3 images from independent brains. PVT, paraventricular nucleus of the thalamus, LHb, medial subnucleus of the lateral habenula.

**Table S1:** LHb Gad2+ cluster markers

**Table S2:** Cluster 2 and cluster I7 DEGs

**Table S3:** GO terms for cluster 2 DEGs

**Table S4:** MDD genes in cluster 2 and cluster I7

**Table S5:** Sex-biased genes in Gad2+ neurons

